# Dynamics of Dentate Gyrus Place Cells and Dentate Spikes During Spatial and Nonspatial Changes in Environments

**DOI:** 10.1101/2025.10.24.684382

**Authors:** Peyton G. Demetrovich, Laura Lee Colgin

## Abstract

The dentate gyrus (DG) is thought to play a key role in the formation of dissociable memory representations for similar contexts. Neurons in the DG receive highly processed spatial and nonspatial sensory information from the medial and lateral entorhinal cortices, respectively. Changes in spatially tuned firing patterns of DG place cells occur after spatial changes to an environment, but the degree to which DG place cells respond to ethologically relevant nonspatial stimuli is largely unknown. Spatial and nonspatial information is thought to be transmitted to the DG during discrete local field potential events called dentate spikes. Here, we tested the extent to which different spatial and nonspatial stimuli modulate place cell firing patterns and dentate spike dynamics. We performed extracellular recordings of DG place cells and local field potentials in rats of both sexes exploring a familiar spatial environment, in which social stimuli and nonsocial odors of varying ethological relevance were presented, and a novel spatial environment. As expected, DG place cells exhibited different firing patterns between familiar and novel environments. Significant changes in firing were not observed, however, with any of the nonspatial stimuli. Surprisingly, the occurrence of dentate spikes associated with lateral entorhinal cortex input increased during exploration of ethologically relevant stimuli, and this increase was greater for social stimuli. Altogether, these results suggest that the DG preferentially responds to social stimuli at the network level, providing novel insights into how spatial and nonspatial information is processed in the DG.

**Significance Statement:** The dentate gyrus (DG) encodes spatial and nonspatial sensory information. Here, we investigated how place cells in the DG respond to changes in spatial and nonspatial cues in familiar and novel environments in rats. We found that DG place cell firing patterns significantly changed in a novel spatial environment but did not significantly change when nonspatial stimuli were presented in a familiar environment. Conversely, discrete dentate spike events reflecting presumed nonspatial inputs from the lateral entorhinal cortex increased during investigation of ethologically relevant nonspatial stimuli. These findings suggest novel mechanisms of nonspatial information processing in the DG.

## Introduction

The hippocampus is a key brain area for spatial and episodic memory formation (Sugar and Moser, 2019). The creation of unique memory representations for different experiences in similar places requires highly correlated inputs to be largely decorrelated, a process known as pattern separation. The dentate gyrus (DG) subregion of the hippocampus is thought to perform pattern separation because it contains a large number of principal neurons that receive a relatively small number of excitatory inputs (Yassa and Stark, 2011; Borzello et al., 2023). Yet, how the principal neurons of the DG accomplish pattern separation during formation of memories for different experiences is not fully understood.

Pattern separation in the DG has been studied in the context of place cells. Place cells are neurons that fire in specific spatial locations called “place fields” (O’Keefe, 1976) and together form updatable cognitive maps of an environment. Transformation of cognitive maps of an environment can occur in response to salient changes in sensory cues or motivational context through a process known as place cell “remapping” (Colgin, Moser, and Moser, 2008). Remapping can involve changes in an individual place cell’s place field location (“global remapping”) or its firing rate (“rate remapping”). A pioneering study of pattern separation in DG place cells revealed that gradual morphing of the shape of an environment induced global remapping in DG place cells (Leutgeb et al., 2007). More recent work determined that both excitatory cell types of the DG, granule cells and mossy cells, can be place cells and can remap when visuo-spatial cues in an environment are changed (Goodsmith et al., 2017, 2019; Senzai and Buzsáki, 2017; Kim et al., 2023; Huang et al., 2024).

Another recent line of research has begun to characterize cue cells, cells with firing selective for sensory cues rather than spatial location (Jung et al., 2019; Woods et al., 2020; Tuncdemir et al., 2022, 2023). Between place cells and cue cells, the DG has the capability to code spatial and nonspatial information in parallel (Fernández-Ruiz et al., 2021; Tuncdemir et al., 2022). This finding raises the possibility that DG place cells will lack responses to nonspatial information such as olfactory cues. On the other hand, spatial and nonspatial information can be integrated through conjunctive encoding in DG neurons (Kim, Jung, and Royer, 2020), suggesting that DG place cells may exhibit remapping to salient olfactory cues. DG granule cells receive olfactory inputs via the lateral entorhinal cortex (LEC; Kerr et al., 2007) and can specifically code odor identity (Woods et al., 2020). Also, place cells in CA2, a hippocampal subregion that receives input from DG granule cells (Kohara et al., 2014; Dudek et al., 2016), selectively remap in response to social but not nonsocial odors (Robson et al., 2025). Together, these findings raise the question of whether DG place cells also remap selectively to social odors.

Dentate spikes are population burst events observed as short duration (< 30 ms), large amplitude (> 1 mV) deflections in the DG local field potential (LFP). Dentate spikes are associated with input from entorhinal projections and modulation of DG cell spiking (Bragin et al., 1995; Dvorak et al., 2021) and have been linked to certain memory operations (Nokia et al., 2017; Farrell et al., 2024; McHugh et al., 2024). Dentate spikes are categorized into two distinct types: DS1 (LEC driven) and DS2 (medial entorhinal cortex (MEC) driven). Given their differential associations with nonspatial LEC and spatial MEC inputs (Hargreaves et al., 2005), DS1s and DS2s are thought to relay nonspatial and spatial information to the DG, respectively. Thus, a simultaneous investigation of place cell activity and dentate spike dynamics may shed light on how the DG codes spatial and nonspatial information. The present study compared neurophysiological recordings of DG place cells and dentate spikes in rats exploring a familiar spatial environment with varied nonspatial stimuli or a novel spatial environment.

## Materials and Methods

### Subjects

Five adult Long-Evans rats (sex: 4 male, 1 female; age: ∼3-8 months at time of surgery; weight: ∼300-700 g) were used in this study. We did not design or power our study to test for potential effects of sex, but our statistical modeling approach accounted for random variability across rats (see *Place cell analyses, Stimulus zone exploration time,* and *Dentate spike analyses* sections, below). Prior to surgery, rats were double or tripled housed with other rats of the same sex on a 12-hour reversed light cycle (lights on: 8 pm-8 am). Rats were acclimated to handling and trained to run clockwise laps on a raised circular track for sweetened cereal pieces. All handling, behavioral training, and recording occurred during the lights off period (8 am-8 pm). After surgery, implanted rats were singly housed in a larger cage with enrichment materials (e.g., wooden blocks and cardboard tubes) and placed directly next to their former cage mates. Behavioral training resumed, and recordings began, as soon as rats recovered from surgery (confirmed by veterinary approval), which occurred as early as the day after surgery. All rats had access to food and water ad libitum. Experiments were conducted according to the guidelines of the United States National Institutes of Health Guide for the Care and Use of Laboratory Animals and under a protocol approved by the University of Texas at Austin Institutional Animal Care and Use Committee.

### Neuropixels probe drive preparation

The silicon probe drive preparation protocol was adapted from Vöröslakos et al. (2021). Before implantation, Neuropixels probes (Jun et al., 2017; Steinmetz et al., 2021) were mounted onto small metal microdrives (“R2Drive”, 3Dneuro, Nijmegen, The Netherlands), which were used to lower probes to target recording areas during surgery (see *Surgery and probe targeting* section below). A layer of epoxy (Loctite Quickset Epoxy) was applied to the Neuropixels headstages to protect the electronics. A Teflon-coated stainless-steel wire with a miniature connector pin (A-M Systems, Sequim, WA) was soldered to the ground connection pad on the Neuropixels flex cable to serve as a ground wire and potential external reference.

### Surgery and probe targeting

Rats were anesthetized with isoflurane vapor (1.5-3%), and 8-9 bone screws were affixed to the skull. Craniotomies were drilled above the right dorsal dentate gyrus (-3.8 mm anteroposterior from bregma and -2.6 mm from the midline), and any obstructing dura was removed from the craniotomy. Two rats (Rats 503 and 504) had an additional craniotomy drilled over the left hippocampus for attempted dual-hemisphere recordings. Due to unforeseen complications with the second probe during both surgeries (e.g., broken probe), the second probe was not implanted in the left hemisphere of either rat. Left-hemisphere craniotomies were sealed with silicone adhesive (“Kwik-Sil”, World Precision Instruments, Sarasota, FL) and covered with cement. One of the bone screws anterior to bregma was connected to the ground wire on the probe. Recordings were monitored during surgery as the probe was stereotaxically lowered into the brain through the craniotomy at a rate of 0.1 mm every 3-5 seconds. Once the probe was lowered to the point where the base of the microdrive contacted the skull, the microdrive was used to advance the probe the remainder of the distance to the target recording sites. The appearance of characteristic DG LFP patterns (i.e., dentate spikes (Bragin et al., 1995; Dvorak et al., 2021) and strong slow gamma (Hsiao et al., 2016; Dvorak et al., 2021)) and cell spiking activity on multiple, deeply located probe channels served to guide probe placement. Probe advancement continued until a full DG LFP depth profile (superior to inferior blade) was visible along the probe. The craniotomy was then sealed with silicone gel (“Dura-Gel”, Cambridge NeuroTech, Cambridge, England), and a layer of Vaseline was applied around the craniotomy and probe shanks to protect them from dental cement. The Vaseline was heated with a cauterizing tool (Bovie Change-A-Tip Deluxe Low Temperature Cautery Kit, MDmaxx, Lakewood, NJ) to facilitate application. An initial layer of dental cement was then used to bond the drive base and bone screws to the skull. A custom 3-D printed conical cap was lowered over the implant and cemented in place to protect the implant from mechanical damage and debris. The Neuropixels headstage was cemented to the inside of the cap, with the probe connected to the headstage through the zero-insertion force connector. This was done to avoid connecting and disconnecting the probe interface on the headstage each day across multiple days of recordings. Instead, the cable interface on the headstage was connected to and disconnected from the data acquisition system each day using the Omnetics connectors on the Neuropixels headstage and interface cable.

### Behavior

Electrophysiological recordings were collected as rats ran clockwise laps on a raised circular track (100 cm diameter) across four 10-minute sessions. Before and after each run session, the rat rested off the track for 10 minutes on a towel-lined stand. After each completed lap, the animal received a sweetened piece of cereal at a specific location of the track that remained constant throughout the experiment. The portion of the circular track spanning 90° around the reward location was designated as the Reward Zone. If a rat turned around mid-lap, a vertical blockade was placed onto the track to prevent the rat from traveling in the counterclockwise direction. The blockade was removed when the animal resumed clockwise travel.

A custom-built stimulus cage (a clear plastic shoebox with holes drilled into the sides) was placed on a separate raised platform directly adjacent to the circular track. During the first session (session A) and fourth session of all conditions, the stimulus cage contained only clean bedding (i.e., standard rat housing bedding). The type of stimulus introduced in the second session (session B), and repeated in the third session, depended on the experimental condition. Sessions A and B of the different experimental conditions are depicted in Fig. 1. For the Empty and Novel Room conditions, the stimulus cage contained only clean bedding in all sessions. In the Social condition, a familiar rat (the implanted rat’s former cage mate) and the familiar rat’s soiled bedding were placed in the stimulus cage in session B. In session B of the Social Odor condition, the stimulus cage contained only familiar rats’ soiled bedding without a familiar rat present. In the Nonsocial Odor condition, a fruit-like odor was presented by adding three drops of hexyl acetate (Sigma-Aldrich Cat# 108154) to the clean bedding of the stimulus cage in session B. In the Fox Odor condition, 3 drops of fox urine (Original Predator Pee, Maine Outdoor Solutions LLC, Hermon, Maine) were introduced to the clean bedding in the stimulus cage in session B. Rats were not discouraged from investigating the stimulus cage unless they spent more than two minutes investigating the cage without completing a single lap. The portion of the track spanning 90° around the stimulus cage was designated as the Stimulus Zone. The remaining 180° of the track that belonged to neither the Stimulus Zone nor the Reward Zone was designated as the Neither Zone.

**Figure 1.**
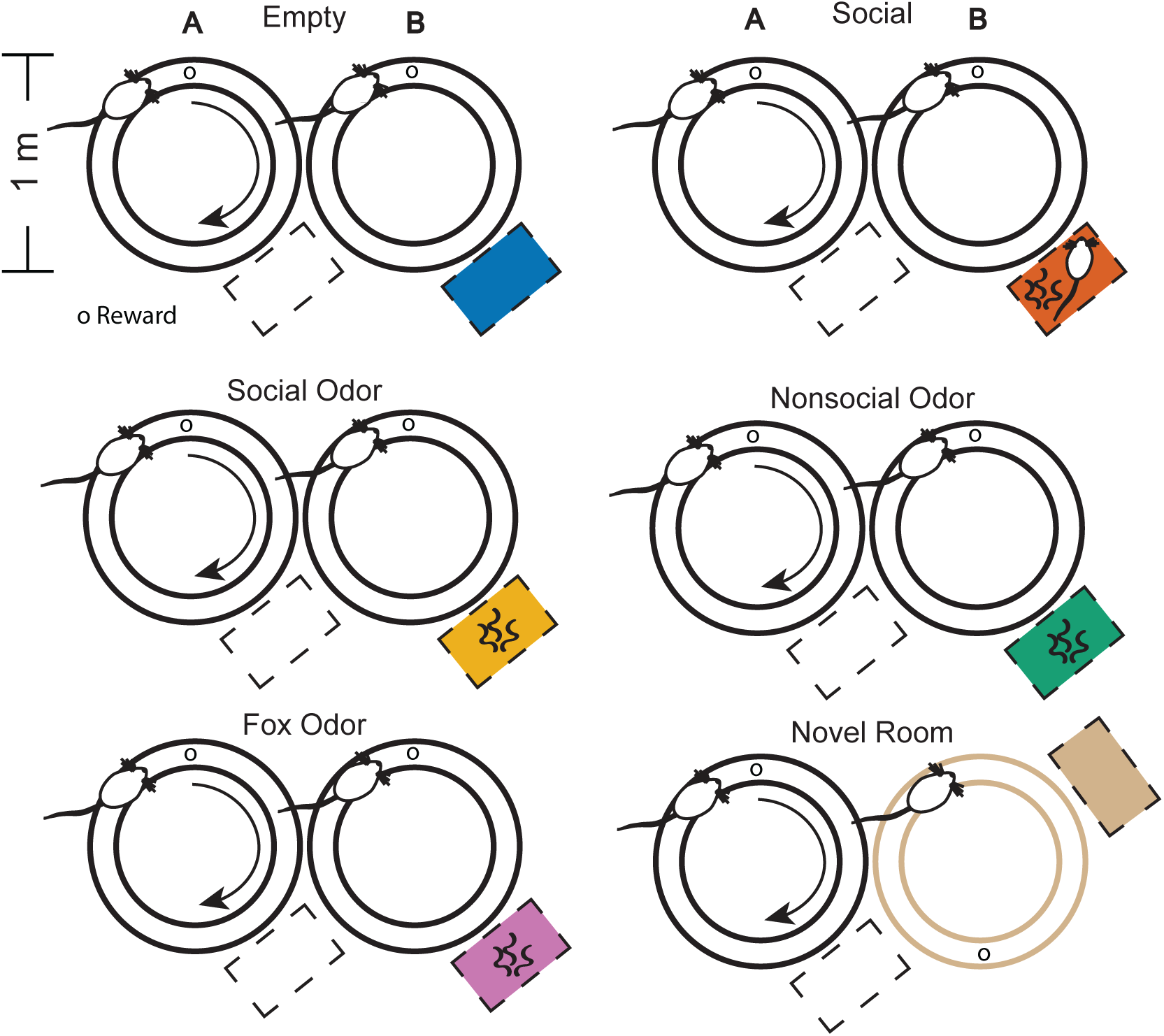
Experimental paradigm. Rats ran clockwise laps on a circular track for food rewards across multiple 10-minute sessions. A stimulus cage was placed on a raised platform adjacent to the circular track. In the first session (A) of all experimental conditions, the stimulus cage contained only clean bedding. Depending on the condition, different stimuli were presented in the stimulus cage in the second session (B). In the Empty condition, the clean bedding remained unchanged. In the Social condition, a familiar rat and its soiled bedding were presented. In the Social Odor condition, the soiled bedding from the home cage of a familiar rat (or familiar rats) was presented. In the Nonsocial Odor condition, a fruit-like odor, hexyl acetate, was added to the clean bedding. In the Fox Odor condition, fox urine was added to the clean bedding. In the Novel Room condition, the implanted rat ran laps in a different spatial environment, a novel room with a different configuration of track and room cues, in session B. The stimulus cage in the novel room contained only clean bedding as in session A.

In the Novel Room condition, recording was stopped after the rest period following session A in the familiar room. The rat was disconnected and moved to a novel recording room that had never been visited before. The rat was then re-connected, and session B recordings resumed in the novel room. The novel room contained a circular track of a different color and track width but identical diameter as in the familiar room. The novel room included the same major landmarks as the familiar room (e.g., stimulus cage, rest stand, acquisition system, computer and monitor, desk, chair, shelving, etc.). However, the spatial relationships among the track, major landmarks, and reward location were different than in the familiar room.

For all experimental conditions, recordings were also collected in the third and fourth sessions (i.e., the sessions in which the environment and stimuli were the same as sessions B and A, respectively). Place maps from these sessions were analyzed and visually inspected to verify spatial selectivity during identification of DG place cells (see *Spike sorting and unit selection* section below). However, these duplicate sessions were excluded from further analyses to simplify the experimental design and corresponding statistical models.

### Histology and shank localization

After the last recording, rats were deeply anesthetized with isoflurane and given a lethal dose of pentobarbital via intraperitoneal injection. Rats were then transcardially perfused with phosphate-buffered saline followed by formalin. Probes were not recovered and remained in the brain for at least 1-hour post-perfusion before brains were collected. This method was selected to enhance the visibility of the Neuropixels shank tracks and facilitate anatomical localization of recording sites. Brains were coronally sectioned at 30 µm, and the sections were stained with cresyl violet (a cell body stain) to verify probe placement in the dorsal dentate gyrus (Fig. S1). Data from shanks that did not penetrate the DG granule cell layer were excluded from analyses.

### Data Acquisition

Recordings were performed using commercially available Neuropixels probes (1.0 or 2.0). Rat 452 was implanted with a 1.0 probe, and all other rats were implanted with 2.0 probes. Signals were collected slightly differently, depending on the probe type. The 1.0 probe amplified, filtered, and digitized the incoming signals into separate action potential (gain = 500, bandwidth = 0.3-10 kHz, sampling rate = 30 kHz) and LFP signals (gain = 250, bandwidth = 0.5-500 Hz, sampling rate = 2.5 kHz). In contrast, the 2.0 probes amplified, filtered, and digitized the incoming signals into a single wideband file (gain = 100, bandwidth = 0.5-10 kHz, sampling rate = 30 kHz). Both probe type’s signal acquisition processes were carried out by the probe’s on-board circuitry. Signals from the probe were transmitted to a Neuropixels PXIe or “OneBox” acquisition system (imec, Leuven, Belgium) via a twisted-pair Neuropixels cable (CBL_1000) that was protected by additional flexible spiral wire wrapping (Altex Computers and Electronics, Austin, TX). SpikeGLX software (https://billkarsh.github.io/SpikeGLX/) was used to run the Neuropixels acquisition system, configure the probe’s channel mapping daily for maximal cell yield, and synchronize the neural and behavioral data streams. Animal position was tracked offline using DeepLabCut position estimation software (Mathis et al., 2018; https://github.com/DeepLabCut) from videos obtained using a single video camera (White Matter e3Vision camera, White Matter LLC, Seattle, WA) located above the circular track. A 30 Hz TTL signal corresponding to frame times of the camera feed was outputted using a camera synchronization device (White Matter e3Vision hub) to an external data acquisition board (NI USB-6259, NI, Austin, TX). This board also received a 1 Hz TTL signal from the Neuropixels PXIe module during recording. The external board was not used for OneBox recordings; instead, camera TTLs were outputted directly to OneBox’s auxiliary input/output board, and the 1 Hz signal was internally generated. Using both TTL signals, SpikeGLX synchronized the data streams to a common reference clock. After recording, the 2.0 probe’s raw 30 kHz wideband data were demultiplexed, artifact cancelled, and filtered into separate spike (0.3-10 kHz; sampling rate = 30 kHz) and LFP (1-500 Hz; sampling rate = 2.5 kHz) files using CatGT software (also available at https://billkarsh.github.io/SpikeGLX/). CatGT was also used to concatenate all spike files across a day’s recording sessions for subsequent spike sorting. The LFP files from seven sessions (Rat 452: Nonsocial Odor, Rat 501: Fox Odor and Novel Room, and Rat 503: Empty Day 1, Social Day 1, Nonsocial Odor Day 1, and Novel Room) were not included in dentate spike analyses due to noise contamination disrupting dentate spike detection and classification.

### Spike sorting and unit selection

Spike sorting was performed using Kilosort4 software (Pachitariu et al., 2024; https://github.com/MouseLand/Kilosort) with default parameters. Next, putative sorted units were evaluated by the automated cell curation software Bombcell (Fabre et al., 2023; https://github.com/Julie-Fabre/bombcell). Bombcell extracts quality metrics such as the number of refractory period violations, spike waveform shape, and the signal-to-noise ratio per unit, which it uses to classify each putative unit as good, multi-units, non-somatic, or noise. Putative units were automatically excluded if 10% or greater of the unit’s spikes violated a 1 millisecond refractory period criterion. Units were then manually curated to check for and perform necessary merge and split operations and verify accurate classification. Manual final unit acceptance generally only included units classified as good, but some multi-units and non-somatic units were accepted due to classification that was perceived as inaccurate during the manual curation step. Cells from the accepted units were then required to meet three additional criteria to be included in further analyses: (1) To be classified as a DG cell, the channel on which the unit displayed its maximal spike waveform had to be within 150 µm of either DG blade’s DS2 sink/source reversal point (Dvorak et al., 2021; Senzai and Buzsáki, 2017). (2) To be classified as an excitatory cell, the unit’s mean firing rate had to be between 0.1 and 10 Hz (Kim et al., 2023). (3) To be identified as a place cell, the unit had to exhibit significant spatial tuning, which was defined as an observed spatial information score (Skaggs et al., 1996) greater than at least 95% of the spatial information scores computed from 100 shuffled rate maps (p < 0.05). Shuffled rate maps were created using the same method as normal rate maps (see *Place cell analyses* section below) except that a given rat’s position coordinates were circularly shifted by at least 30 s. Cells that met all 3 additional criteria were classified as DG place cells.

### Place cell analyses

One-dimensional firing rate maps were constructed using previously published methods with similar parameters (Hwaun and Colgin, 2019). First, each unit’s spikes were binned into 6° position bins across the full 360° of the circular track using the rat’s position at the spike time. For each position bin, the total number of binned spikes per unit was divided by the amount of time the rat spent in that bin. Only spikes that occurred while the rat was moving at least 5 cm/s were included. Raw rate maps were subsequently smoothed by convolution with a one-dimensional Gaussian kernel (standard deviation = 15°). Smoothed and linearized rate maps from sessions A and B for all cells included in analyses are shown in Fig. 2.

**Figure 2.**
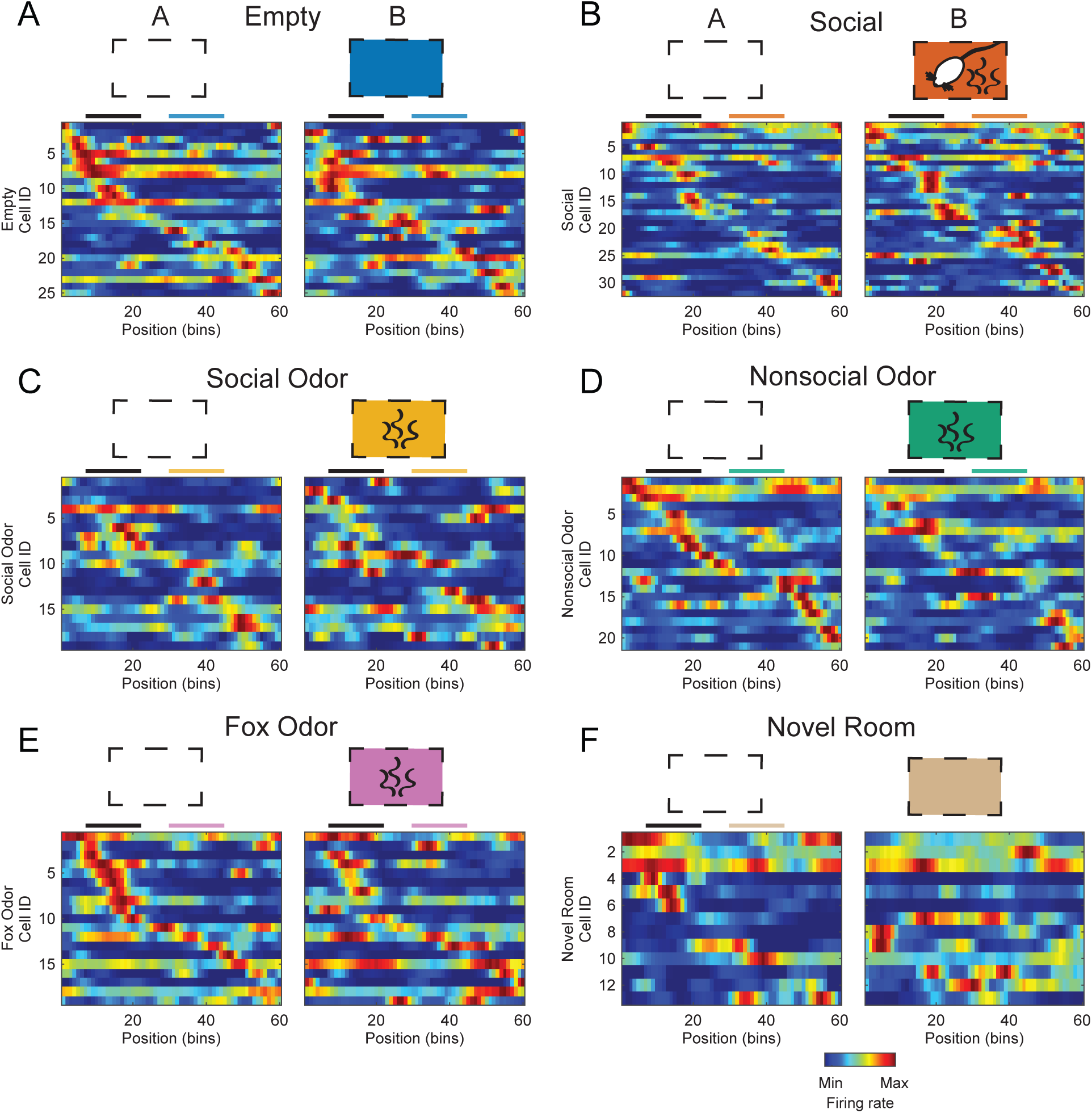
One-dimensional firing rate maps for all DG place cells included in analyses (recorded across rats and days and plotted per condition and session). Each spatial bin corresponds to 6° (out of 360° of positions along the circular track). Rate maps are shown sorted by the location of maximal firing on the track in session A for each condition and scaled to each cell’s maximal firing rate across sessions. Above the rate maps, the black line denotes the position bins of the track where the rats received a food reward after each lap (Reward Zone), and the colored line denotes the bins closest to the stimulus cage (Stimulus Zone).

Global remapping of place cells was assessed by calculating the spatial correlation coefficient (i.e., Pearson correlation) between the rate maps from sessions A and B for a given cell (Fig. 3A). To determine if spatial correlations significantly differed between conditions, we employed a linear mixed model (LMM) approach using the MATLAB Statistics and Machine Learning Toolbox (The MathWorks, Inc.) function *fitlme*. Spatial correlation values were non-normally distributed and bounded [-1,1], so the Fisher r-to-z transformation (z = atanh(r)) was applied to all spatial correlation values prior to fitting the model. Transformed spatial correlation was specified as the response variable (i.e., dependent variable) and was modeled as a function of condition as a fixed effect and rat as a random effect. The transformed spatial correlation values of the Empty condition (i.e., the control group) were used as the reference level for dummy variable coding. A main effect of condition was assessed within the fitted LMM using the MATLAB LMM object function *anova*. The estimated marginal means and 95% confidence intervals for each condition’s spatial correlation values were calculated using the MATLAB LMM object function *predict* and are shown in Fig. 3B. One-dimensional k-means clustering (number of clusters = 2) was performed on the Empty control condition’s raw spatial correlation values to identify clusters of stable (non-remapping) and unstable (remapping) cells. Cluster centroid locations were then averaged to obtain a stability threshold separating the two clusters of cells.

**Figure 3.**
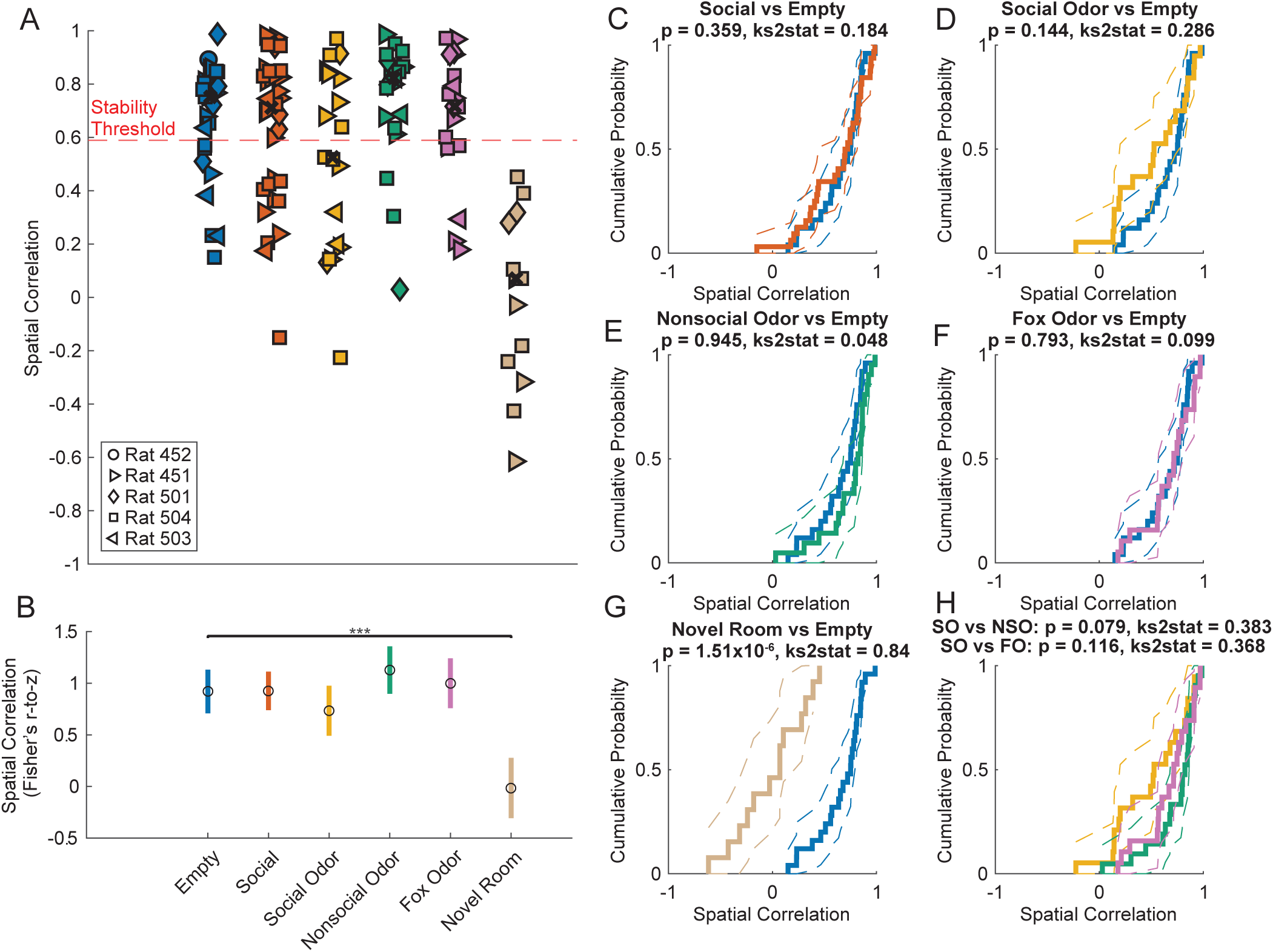
Propensity of global remapping in DG place cells compared across conditions. A. Spatial correlation values between session A and B rate maps for all DG place cells across conditions. Each marker represents a single cell, and the marker shape reflects the rat from which the cell was recorded. Black x’s show each condition’s median spatial correlation value. The horizontal dashed line denotes the remapping threshold used to separate globally remapping and non-globally remapping cells (see Materials and Methods). B. Mean Fisher-transformed spatial correlation values estimated from a LMM are shown across conditions. Open circles denote marginal means, and colored lines represent associated 95% confidence intervals. C-G. Cumulative distribution functions of spatial correlation values for each condition are shown. Dashed lines represent the 95% confidence intervals for each distribution. The control (i.e., Empty) condition’s distribution is replicated in blue in each plot for ease of comparison. H. Comparison of cumulative distribution functions for Social Odor (SO), Nonsocial Odor (NSO), and Fox Odor (FO) conditions. Two-sample Kolmogorov-Smirnov test statistics (ks2stat) and uncorrected p-values are reported above each plot for the corresponding distributions. Note that significant global remapping was only observed in the Novel Room condition (* indicates p ≤ 0.05, ** indicates p ≤ 0.01, *** indicates p ≤ 0.001).

Rate remapping was assessed by analyzing the difference in the firing rates from sessions A and B for cells with spatial correlation values above the stability threshold only (Fig. 4A). Firing rate changes were calculated as the natural logarithm of the ratio of the mean firing rate in session B to the mean firing rate in session A. This metric, henceforth referred to as the log rate ratio, is signed (that is, positive values indicate increased firing rate in session B while negative values indicate decreased rates), symmetric about 0 (no change in firing rate), unbounded, and respects the approximately lognormal distribution of hippocampal firing rates (Buzsáki and Mizuseki, 2014). Mean firing rates were extracted from each cell’s session-specific rate map. To investigate whether the log rate ratios differed between conditions, we used the same LMM approach as described above for spatial correlations, except with log rate ratio specified as the response variable. The estimated means and 95% confidence intervals for each condition’s values are shown in Fig. 4B.

**Figure 4.**
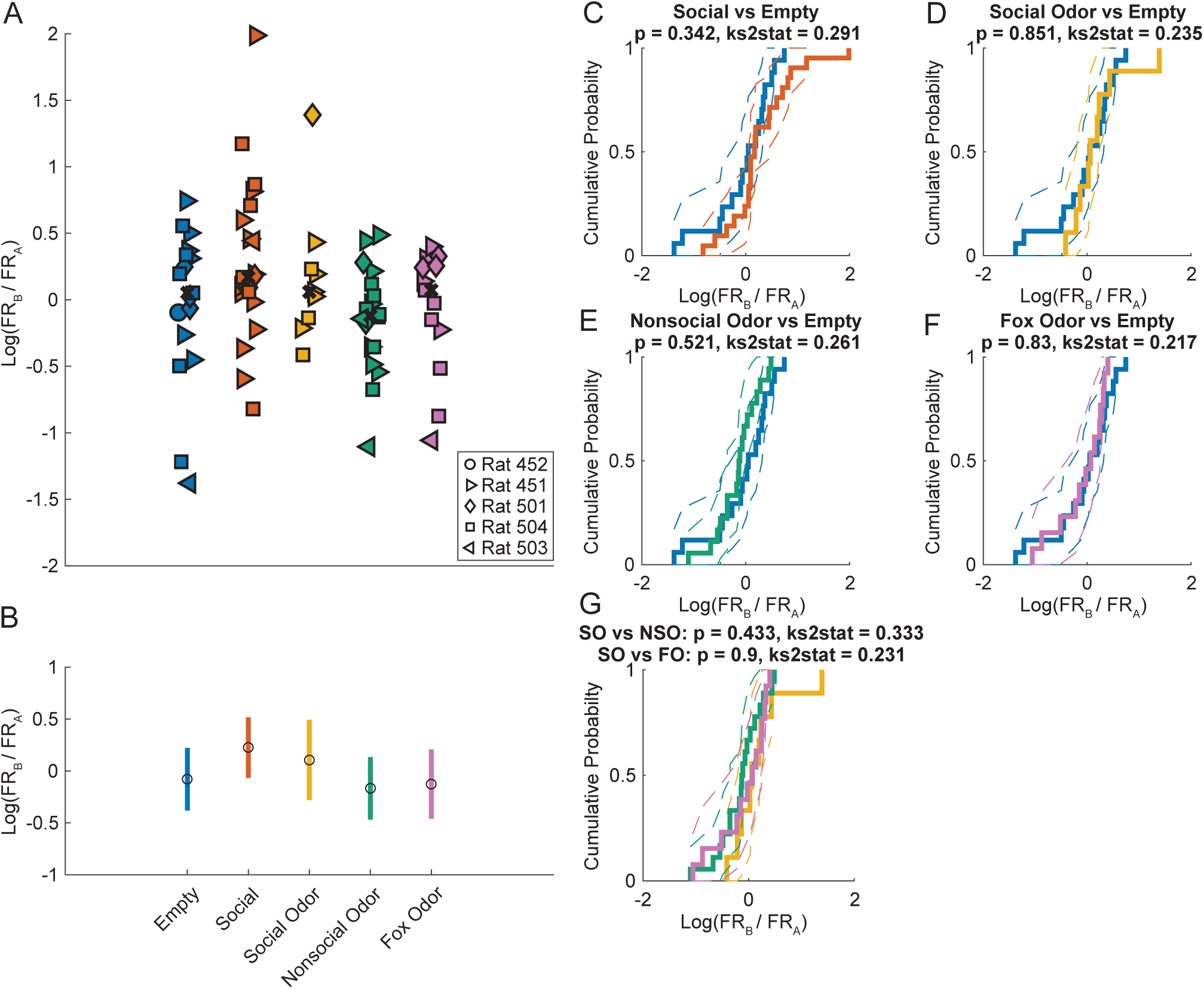
Propensity of rate remapping in spatially stable DG place cells compared across conditions. A. Log firing rate ratios between the DG place cell mean firing rates from session A (FR_A_) and B (FR_B_) rate maps across conditions except for Novel Room (as all cells were classified as unstable between the familiar and novel rooms). Each marker represents a single cell, and the marker shape indicates the rat from which the cell was recorded. Black x’s show each condition’s median log rate ratio. B. Estimated means of log rate ratios from a LMM are shown across conditions. Open circles represent marginal means, and colored lines represent 95% confidence intervals. C-F. Cumulative distribution functions of log rate ratios for each condition. Dashed lines represent the 95% confidence intervals for each distribution. The control (i.e., Empty) condition’s distribution is replicated in blue in each plot for ease of comparison. G. Comparison of cumulative distribution functions for Social Odor (SO), Nonsocial Odor (NSO), and Fox Odor (FO) conditions. Two-sample Kolmogorov-Smirnov test statistics (ks2stat) and uncorrected p-values are reported above each plot for the corresponding distributions. Note a lack of significant rate remapping for any condition.

Cumulative distribution functions with 95% confidence intervals for non-transformed spatial correlation values and log rate ratios for each condition were computed using the MATLAB function *ecdf.* A two-sample, one-tailed Kolmogorov-Smirnov test (MATLAB function *kstest2*) was used to test for a smaller spatial correlation cumulative distribution function (i.e., greater values), and thus a lower degree of remapping, in the Empty control condition compared to the distributions in each of the remaining conditions (Figs. 3C-G). A two-tailed Kolmogorov-Smirnov test was used instead when a specific directional shift in the distribution was not hypothesized a priori (Fig. 3H, 4C-G).

### Stimulus zone exploration time

We collected the time rats spent in each 6° position bin along the circular track during the entire 10 minutes of all recording sessions to create exploration time heatmaps. These heatmaps were averaged across days and rats per condition and session to generate the average exploration time maps shown in Fig. 5A. Exploration times for the bins corresponding to the 90° Stimulus Zone were summed for sessions A and B separately to get a Stimulus Zone exploration time. To compare Stimulus Zone exploration time across conditions and sessions, a generalized LMM (GLMM) was employed using the MATLAB Statistics and Machine Learning Toolbox function *fitglme* with Stimulus Zone exploration time as the response variable, condition and session as fixed effects, rat as a random effect, and an additional random effect of day nested within rat (as repeated presentations of the same condition may influence behavior). Dummy variables were constructed using effects coding in this model. Individual Stimulus Zone exploration times and 95% confidence intervals across conditions and sessions are shown in Fig. 5B.

**Figure 5.**
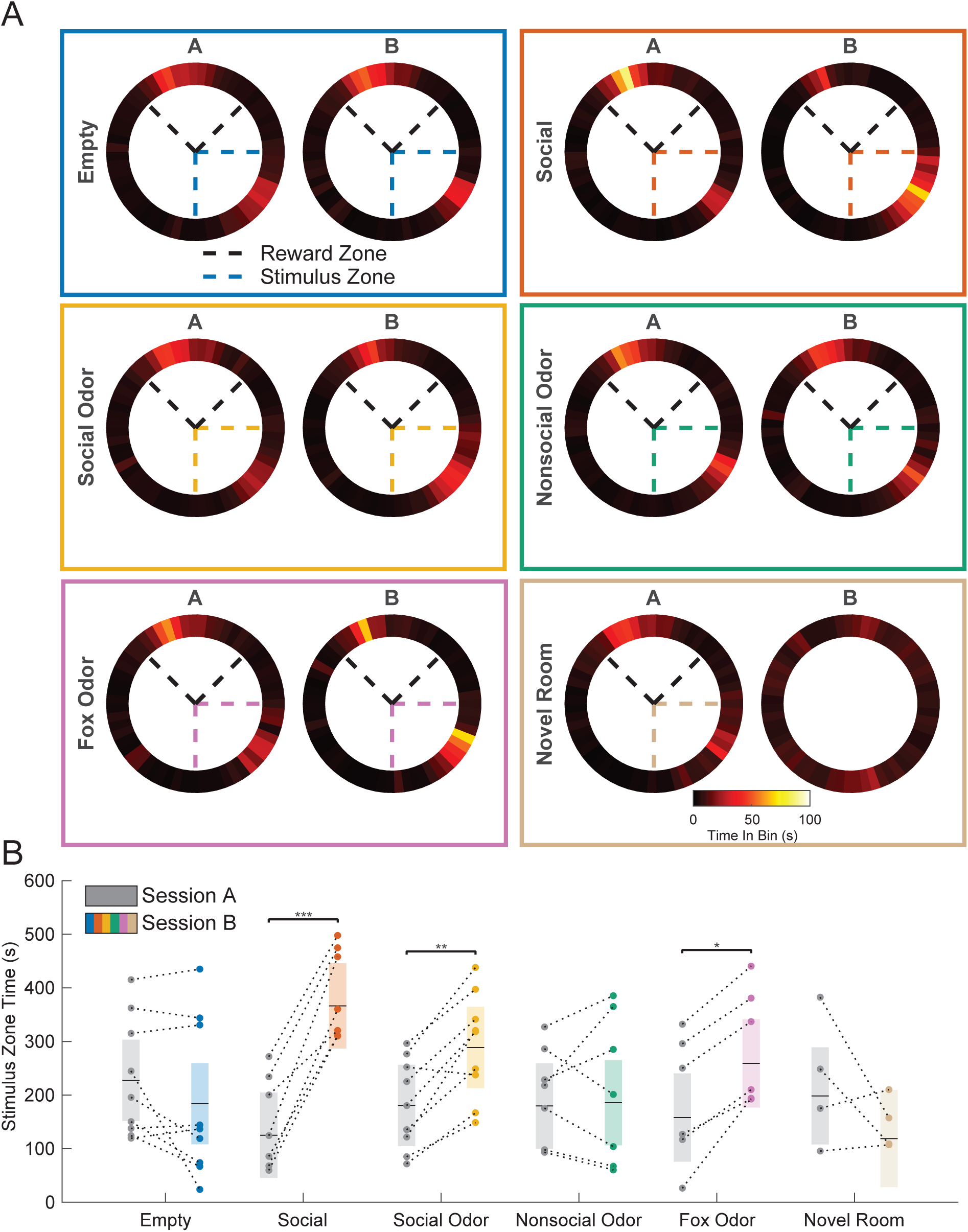
Rats spent more time exploring ethologically relevant stimuli. A. Heatmaps representing the average time rats spent in each track position bin in sessions A and B of each condition. Dashed lines represent the 90° Reward (Black) and Stimulus (Color) Zones. The zones for session B of the Novel Room condition are not shown because their locations were not held constant across recordings (i.e., recordings across sessions were collected in different rooms in the Novel Room condition). B. Individual Stimulus Zone times for each condition and session. Shaded boxes represent the corresponding 95% confidence intervals, and horizontal black lines denote the estimated means for each condition-session pair. Note that rats spent more time in the Stimulus Zone in session B than in session A of the Social, Social Odor, and Fox Odor conditions.

### Dentate spike analyses

Dentate spikes were automatically detected per recording session using methodology adapted from a prior study of dentate spikes in mice (Farrell et al., 2024). First, the dentate LFP channel showing the highest amplitude putative dentate spikes was identified by visual inspection, z-scored, and filtered between 5-100 Hz. Peaks in the filtered signal that exceeded a standard deviation threshold were designated as dentate spikes. To identify a standard deviation threshold in the filtered signal that best detected dentate spikes in our dataset, five different thresholds (2.5, 3.0, 3.5, 4.0, and 4.5) were tested on two different held-out recording sessions. Two blind raters manually identified dentate spikes from these recording sessions to provide a ground truth. An F1 score, the harmonic mean of precision (true positives / (true positives + false positives)) and recall (true positives / (true positives + false negatives)), was computed for each automated and manual detection pair to create a performance curve. Performance curves were averaged across manual scorers, and the threshold value with the consistently highest F1 score was chosen (Fig. S2A). Dentate spikes were then detected using the ground truth-validated standard deviation threshold of 4.0 for all experimental conditions. Example current source density (CSD) depth profiles around the peaks of dentate spikes detected by different methods are shown in Fig. S2B. CSD profiles were obtained by transforming the LFP depth profiles around dentate spike times (±50 ms) using the spline inverse CSD method within the CSDplotter MATLAB toolbox (Pettersen et al., 2006; https://github.com/espenhgn/CSDplotter).

Detected dentate spikes were then split into two separate types (putative DS1/L or DS2/M) based on k-means clustering (number of clusters = 2) on the first two principal components of their z-scored CSD depth profiles (Fig. S2C-D). The average CSD profile for each cluster was computed, and the locations of the sink/source reversals were compared. A cluster was classified as “DS1” if its reversal location was more dorsal than the other cluster (only the CSD profile from the superior blade was used for this analysis). The remaining cluster was then classified as “DS2”. Each cluster’s CSD profile was then visually inspected to verify accurate classification (Fig. S2E). Example dentate spikes classified using this approach are shown in Fig. S3.

Dentate spike rates per type were analyzed in two ways. First, dentate spikes that occurred during times when theta (5-10 Hz) rhythms occurred were excluded. Theta rhythm epochs were identified from the channel with maximal theta power. The channel with maximal theta power was identified for each recording session by first finding the longest bout of time within each session when the rat ran faster than 5 cm/s. This epoch presumably contained prominent and sustained theta. Next, for each channel, the time-frequency representation of power in the theta band during this running epoch was computed using complex Morlet wavelet convolution. Power was then averaged over time per channel, and the channel with maximal theta power was selected for theta detection. Next, the entire LFP from the channel with maximal theta was filtered in the theta band, and the theta amplitude envelope was estimated from the absolute value of the Hilbert transform of the theta-filtered signal. This envelope was z-score normalized, and epochs in the z-scored signal that exceeded a 2-standard deviation threshold for at least 3 average theta cycle lengths were identified as theta epochs. The remainder of recordings were classified as non-theta epochs. The number of DS1 and DS2 events detected in non-theta epochs were then counted per condition and session. To analyze dentate spike rate for different dentate spike types across conditions and sessions, a Poisson GLMM was employed using the MATLAB Statistics and Machine Learning Toolbox function *fitglme*, with a log link function, log(LFP duration) offset variable, and dummy variables constructed using effects coding. The dentate spike count was modeled as a function of the fixed effects of condition, session, and dentate spike type, the interaction effects between the fixed effects, and random effects of rat and day (nested within rat). Individual DS1 and DS2 rates and 95% confidence intervals per condition and session are shown in Figs. 6B and 7B, respectively.

**Figure 6.**
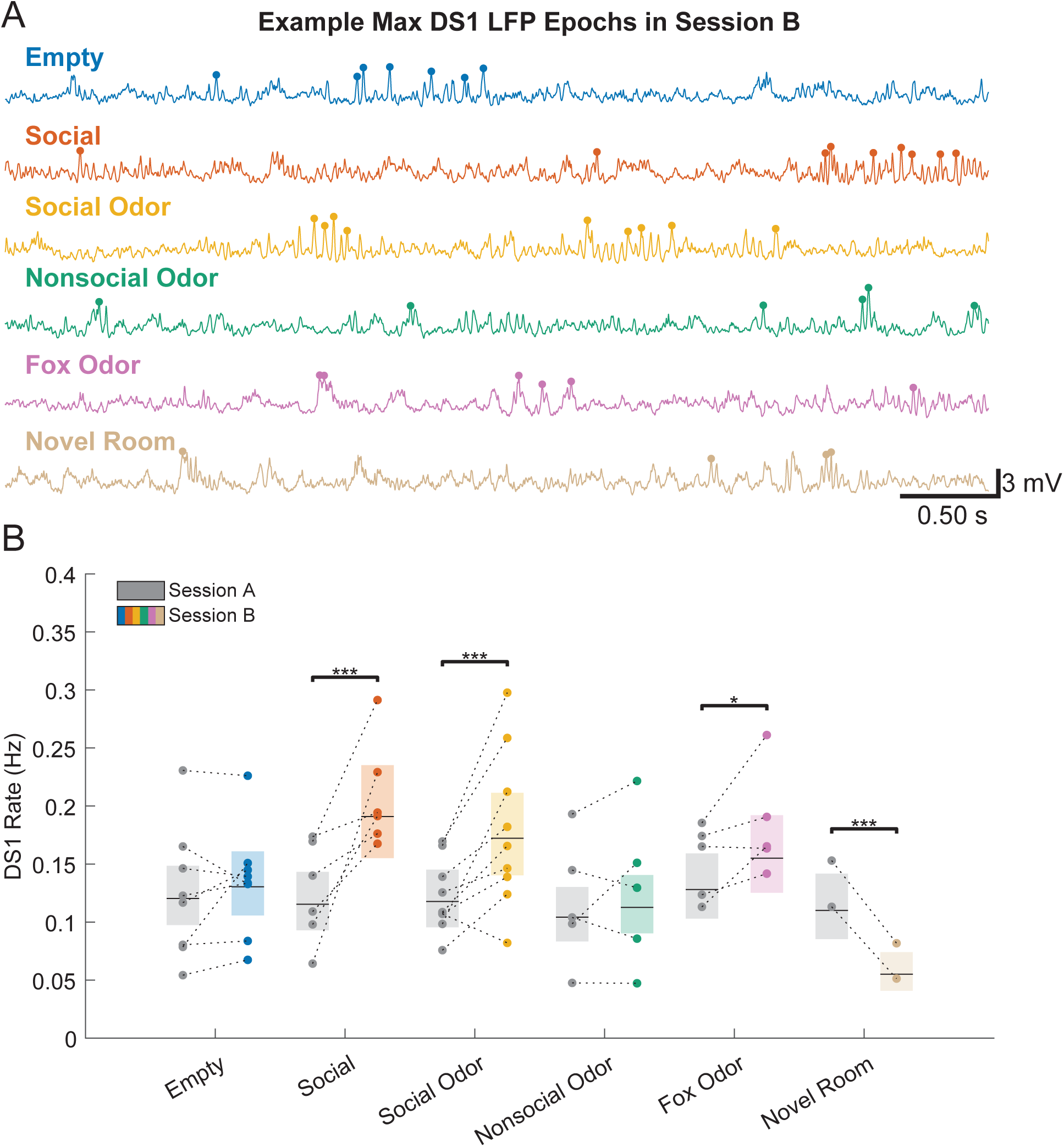
DS1 rates increased during presentation of ethologically relevant stimuli. A. Example LFP recordings containing DS1 events are shown for one example rat. Shown are the five-second epochs that exhibited maximal DS1 rates per condition in session B. The colored dots above positive-going LFP deflections indicate detected DS1s. Note the apparent increase in DS1 occurrence in the social stimuli conditions. B. DS1 rates in sessions A and B are shown across conditions and rats. Each circle represents the DS1 rate from a single session. Shaded boxes represent the 95% confidence intervals, and horizontal black lines denote the estimated means for each condition-session pair. Note that DS1 rates increased in the presence of ethologically relevant odors but decreased in a novel room.

**Figure 7.**
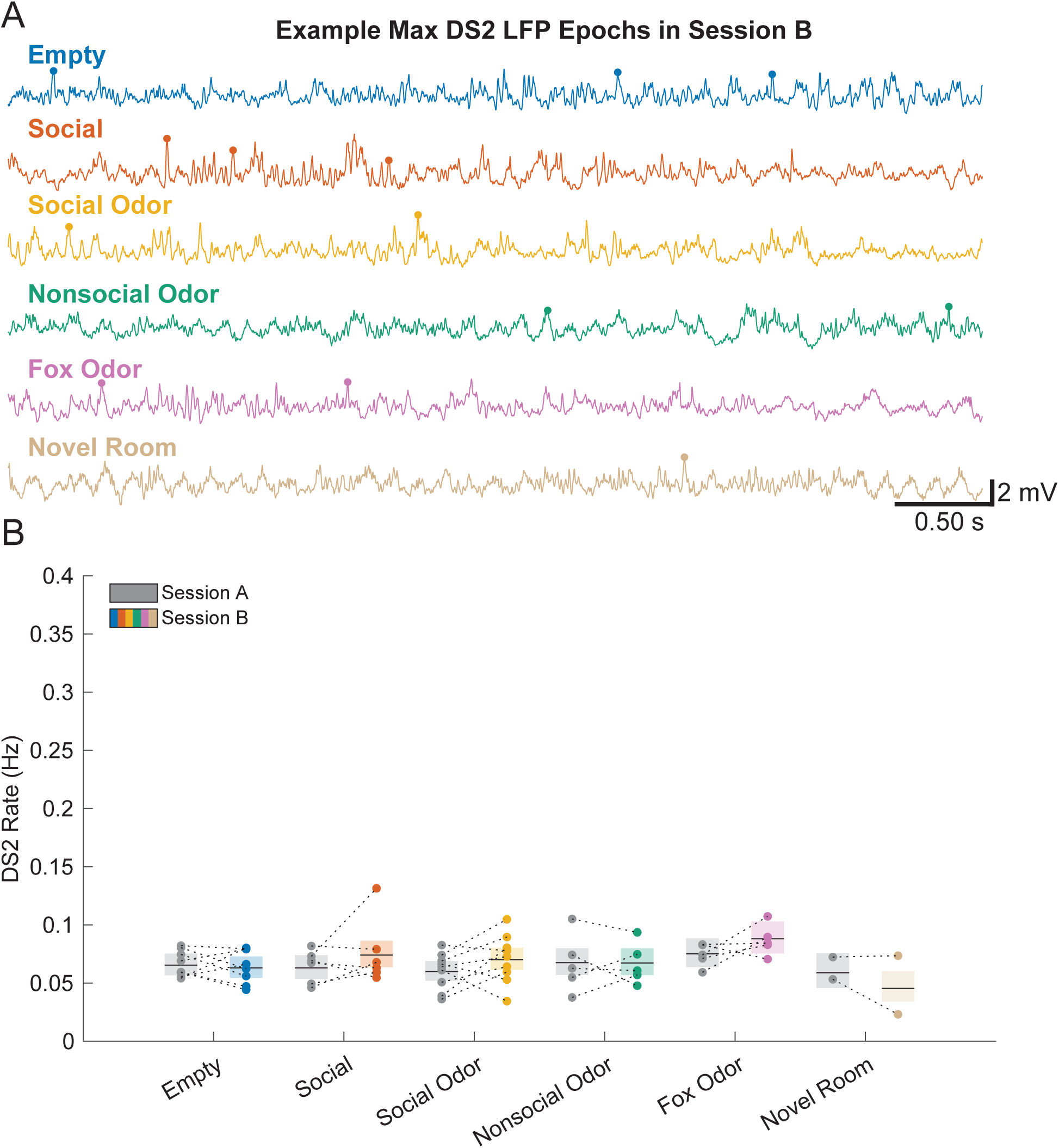
DS2 rates did not change significantly during presentation of nonspatial stimuli or exploration of a novel environment. A. LFP recordings showing the five-second epochs associated with the highest DS2 rate per condition for an example rat. Colored dots indicate detected DS2 events. Note the overall low rate of DS2s, and that no change in DS2 rate is apparent for any condition. B. DS2 rates in sessions A and B are shown across conditions and rats. Each circle represents the DS2 rate from a single session. Shaded boxes represent the 95% confidence intervals and horizontal black lines denote the estimated means for each condition-session pair.

For the DS1 rates shown across position bins in Fig. 8A-F, a “DS1 rate map” was created per session in a similar manner to the place cell firing rate maps described previously. Instead of spike times, however, DS1 times were binned according to the position of the rat at each event time. The number of events in each bin was then divided by the total amount of time the rat spent in that bin. No minimum velocity criterion was enforced as dentate spikes occur most frequently during immobility (Bragin et al., 1995). Raw DS1 rate maps were smoothed with a Gaussian kernel (standard deviation = 15°), and zone-specific rates were extracted from the smoothed rate maps by selecting the bins that corresponded to the different zones on the track and then averaged. These zone-specific mean rates were modeled, again using a GLMM approach, with condition, session, and zone type (i.e., Stimulus Zone, Reward Zone, or Neither Zone), as fixed effects along with the corresponding interaction effects between these fixed factors. Rat and day (nested within rat) were again included as random effects to account for random variability across rats and multiple recording days within rats. Dummy variables were constructed using effects coding. Average DS1 rates and 95% confidence intervals within each zone across conditions and sessions are shown in Fig. 8G-I.

**Figure 8.**
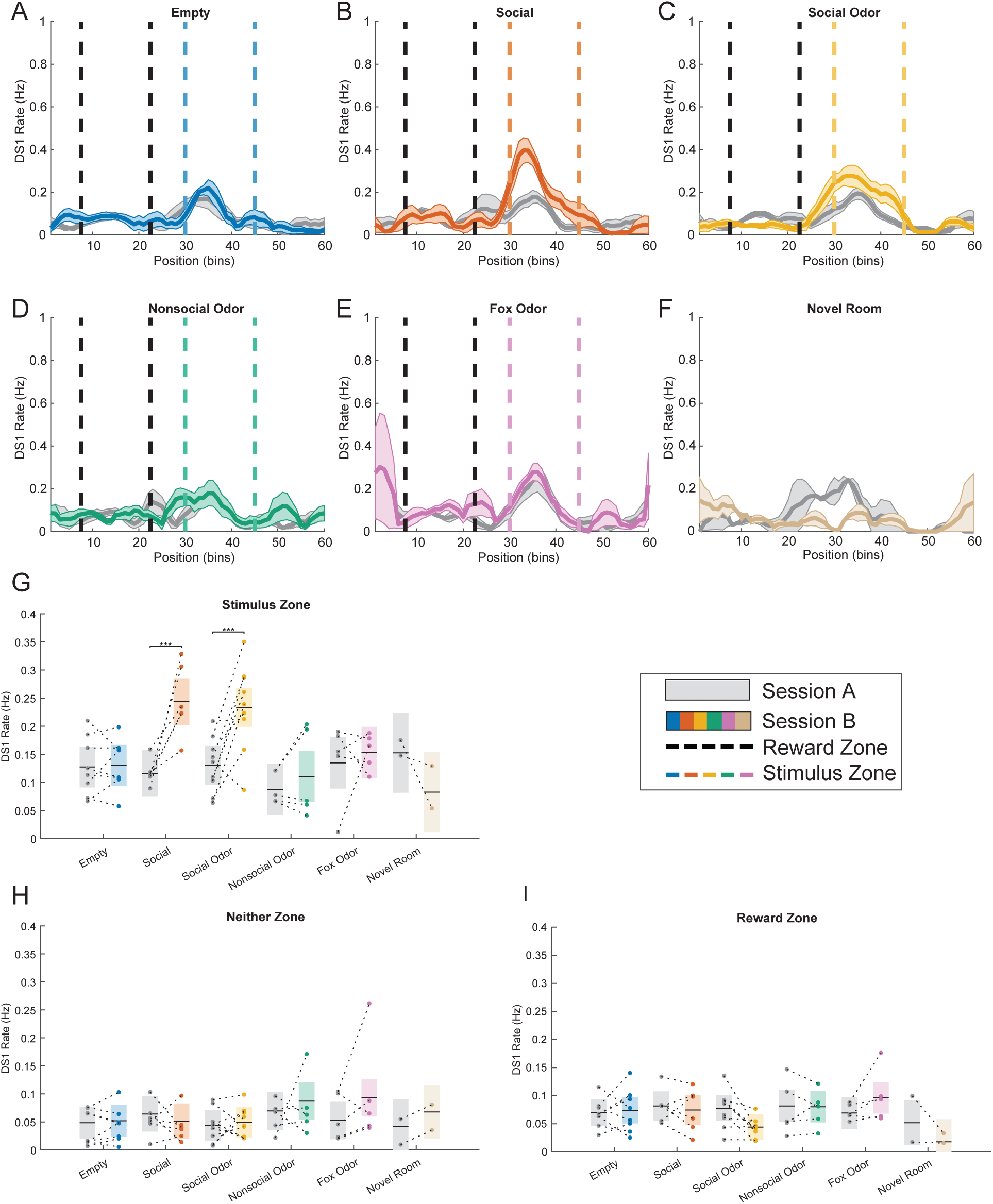
DS1 rates specifically increased during exploration of social stimuli. A-F. DS1 rates across locations on the circular track are shown for all conditions. Shown are the mean ± standard error of DS1 rates across position bins (averaged over repeat condition days and rats) in session A (gray) and session B (colors). Dashed black lines denote the 90° Reward Zone on the track (i.e., the location where the animal received a food reward at the end of each lap), and dashed colored lines denote the 90° Stimulus Zone (i.e., where the stimulus cage was located). Zones are not depicted for the Novel Room condition as their locations were not constant across recordings in different rooms. Each position bin represents 6° on the circular track, such that 60 position bins show the entirety of the 360° track. Note that increases in DS1 rates are apparent within the Stimulus Zone during session B for the Social and Social Odor conditions only. G-I. DS1 rates per condition, session, and track zone. Each circle represents the DS1 rate averaged within the corresponding zone per condition-session pair. Shaded boxes represent the 95% confidence intervals, and horizontal black lines denote the estimated means for each condition-session pair.

DS1 and DS2 peri-event time histograms (PETHs; Figs. 9A and S4A) were constructed by binning DG place cell spikes into 0.01 second bins across a 0.2 second window centered around DS1 peaks (±0.1 s). Binned place cell spike counts were averaged across DS events and divided by the length of the time bin to estimate firing rates (in Hz). Each cell’s average PETH was then smoothed across time with a 5-element long Gaussian-weighted moving average. To estimate a firing rate modulation value per cell for a given condition and session, the average firing rate within the baseline window of -0.1 s to -0.06 s from the dentate spike peak was subtracted from the average firing rate within a window of ±0.02 s around the dentate spike peak. These DS-associated firing rate modulation values were modeled with a GLMM as a function of the fixed effects of condition, session, and dentate spike type, the corresponding interaction effects, and the random effects of rat, cell, and day (with cell and day both nested within rat). Dummy variables were again constructed using effects coding. Firing rate modulation values and 95% confidence intervals for DS1s and DS2s are shown in Figs. 9B and S4B, respectively.

**Figure 9.**
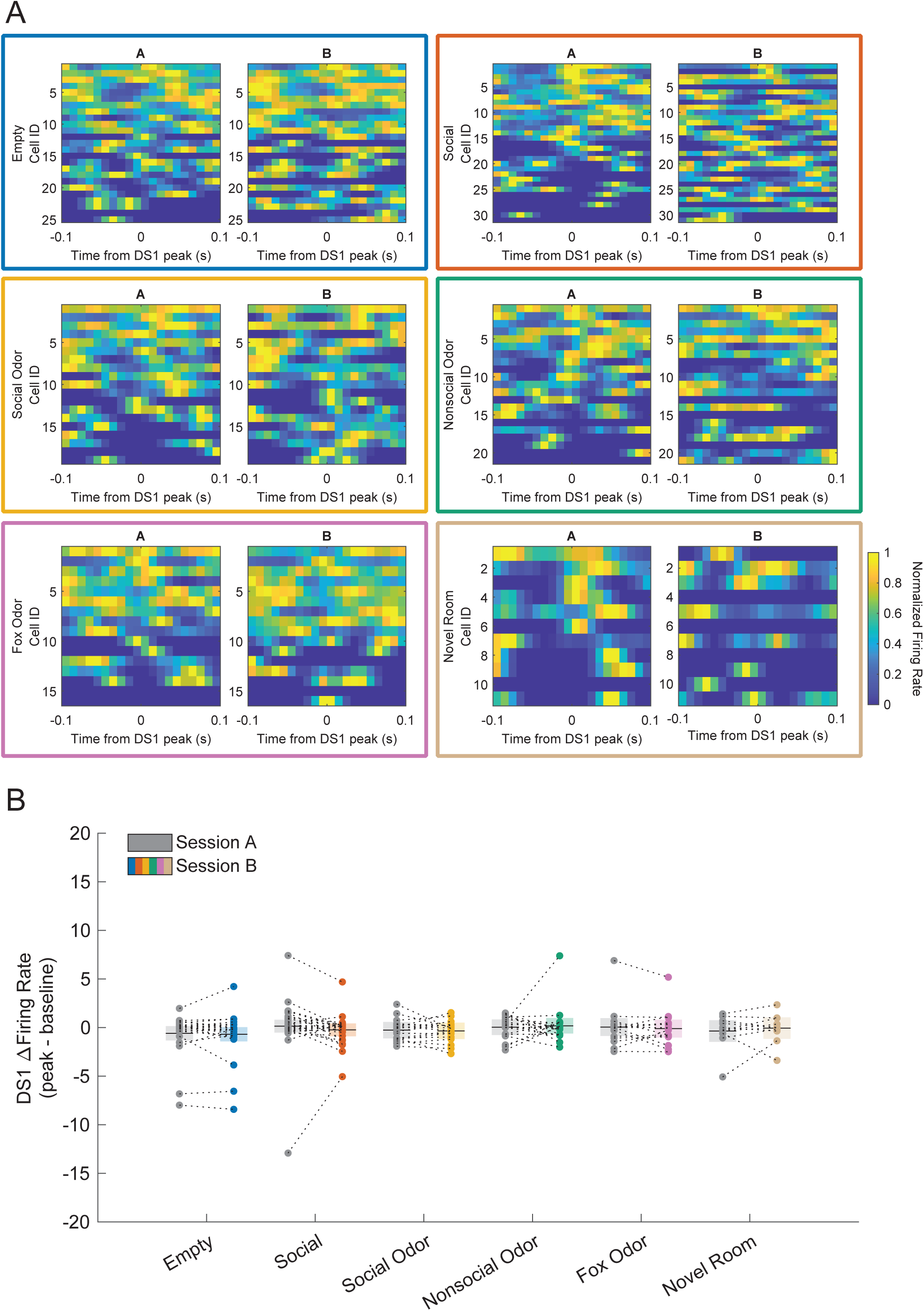
DG place cell firing was not consistently modulated by DS1s. A. Mean peri-event time histograms (PETHs; bin size = 0.01 s) of DG place cell firing rates, constructed around DS1 peaks in sessions A and B, are shown for each condition. Cells are sorted by the maximum firing rate at time 0 in session A per condition. Note that DS1-associated firing was low regardless of session and condition. B. Firing rate modulation values per cell across sessions and conditions. Firing rate modulation was calculated as the average rate from the PETH in a window centered around the DS1 peak (±0.02 s) minus the average firing rate in a baseline window (-0.1 s to -0.06 s before the DS1 peak). Shaded boxes represent the 95% confidence intervals, and horizontal black lines denote the estimated means for each condition-session pair.

### Statistics, data, and code availability

GLMMs or LMMs and two-sample Kolmogorov-Smirnov tests were used to analyze results, as described above for each measure. Statistical tests on each (G)LMM were computed as Wald F-tests or converted T-tests, as reported. Converted T-tests were necessary for one-tailed tests that were used when effects in a specific direction were hypothesized a priori. Post-hoc comparisons were computed only when a significant main effect (for (G)LMMs with one fixed effect) or interaction effect (for (G)LMMs with two or more fixed effects) was observed. All reported P-values were Bonferroni corrected unless otherwise stated. All data and statistical analyses were performed using MATLAB scripts, which are hosted at https://github.com/ColginLab/Demetrovich_Colgin_2026. Scripts were written using custom, built-in, and/or toolbox-specific functions. Certain custom functions were used from our lab’s code base (https://github.com/ColginLab/ColginLabCode). Data will be made available upon reasonable request.

## Results

### DG place cell responses to social and nonsocial olfactory stimuli

Our earlier studies of CA2 place cells showed significant changes in place field locations (i.e., global remapping) when social stimuli were presented (Alexander et al., 2016; Robson et al., 2025). Also, a prior study in mice reported that DG place cells globally remap in response to changes in visual and tactile features of an environment (Kim et al., 2023). Thus, we investigated the extent to which DG place cells exhibit global remapping in response to presentation of social and nonsocial olfactory cues in a familiar environment. To assess global remapping in DG place cells, we calculated spatial correlations between DG place cell firing rate maps from sessions A and B for each condition (Figs. 1-3) and observed a significant main effect of condition (Fig. 3A-B, F(5,123) = 8.69, p = 4.46 x 10^-7^). We hypothesized that spatial correlation values would be significantly lower in all experimental conditions than in the Empty condition.

However, we instead found that spatial correlation values were not significantly lower in the Social (post-hoc one-tailed T-test for Empty vs. Social: t(123) = 0.04, uncorrected p = 0.52), Social Odor (post-hoc one-tailed test for Empty vs. Social Odor: t(123) = -1.15, uncorrected p = 0.13), Nonsocial Odor (post-hoc one-tailed test for Empty vs. Nonsocial Odor: t(123) = 1.32, uncorrected p = 0.90), and Fox Odor (post-hoc one-tailed test for Empty vs Fox Odor: t(123) = 0.49, uncorrected p = 0.69) conditions compared to the Empty control condition. Spatial correlation values in the Social Odor condition were slightly lower than those observed in the Nonsocial Odor condition (post-hoc two-tailed Wald F-test for Social Odor vs. Nonsocial Odor: F(1,123) = 5.45, uncorrected p = 0.02). On the other hand, spatial correlation values in the Social Odor condition did not differ from not Fox Odor values (Social Odor vs. Fox Odor: F(1,123) = 2.37, uncorrected p = 0.13). None of these p-values were significant following corrections for multiple comparisons (Bonferroni-corrected p-value of 0.0143 for 1-tailed tests and 0.0071 for 2-tailed tests). These findings suggest that significant global remapping was not observed in DG place cells during presentation of social stimuli and nonsocial olfactory stimuli in a familiar environment. Significant global remapping was observed, however, during exploration of the novel room environment (Fig. 2F, Fig. 3A-B; post-hoc one-tailed test for Empty vs. Novel Room: t(123) = -5.13, p = 3.80 x 10^-6^), indicating that our population of DG place cells created distinct maps to represent different spatial environments, as expected based on prior studies (Goodsmith et al., 2017; Senzai and Buzsáki, 2017; Allegra et al., 2020). We next compared whether distributions of spatial correlation values differed between conditions (Fig. 3C-H). Only the distributions of spatial correlation values between the Empty and Novel Room conditions were significantly different (two-sample, one-tailed Kolmogorov-Smirnov test, ks2stat = 0.84, p = 1.05 x 10^-5^).

DG place cells also exhibit rate remapping across different environments (Kim et al., 2023). Rate remapping is thought to be driven by LEC inputs carrying nonspatial contextual information (Lu et al., 2013). Here, we assessed the degree of rate remapping using a log rate ratio measure in DG place cells with stable spatial firing (i.e., non-globally remapping cells; see *Place cell analyses* section of Materials and Methods) during presentation of social and nonsocial olfactory stimuli (Figs. 1-2, 4). Since all cells globally remapped in the Novel Room condition, the Novel Room condition was excluded from this rate remapping analysis. Log rate ratios did not significantly differ across conditions (Fig. 4A-B; no significant main effect of condition: F(4,73) = 1.97, p = 0.11). Similarly, distributions of log rate ratios did not differ across conditions (Fig. 4C-G). These results indicate that stable DG place cells do not significantly change their firing rates in response to presentation of social and nonsocial olfactory stimuli.

### Rats explored ethologically relevant stimuli

A potential explanation for the lack of remapping observed in our DG place cell population in response to nonspatial cues could be that the stimuli used were not salient enough to induce stimulus exploration. To assess this possibility, we investigated the amount of time that rats spent within the 90° Stimulus Zone in sessions A and B for all conditions (Fig. 5A). We observed a significant interaction effect between condition and session on time spent in the Stimulus Zone (F(5,72) = 11.52, p = 3.43 x 10^-8^). Compared to Stimulus Zone exploration time in session A, rats spent significantly more time within the Stimulus Zone in session B in the Social (two-tailed Wald F-test for Social-Session A vs. Social-Session B: F(1,72) = 51.55, p = 3.09 x 10^-9^), Social Odor (Social Odor-Session A vs. Social Odor-Session B: F(1,72) = 13.31, p = 2.98 x 10^-3^), and Fox Odor (Fox Odor-Session A vs. Fox Odor-Session B: F(1,72) = 7.76, p = 0.04) conditions. Stimulus Zone exploration times were not significantly different between sessions A and B for the Empty, Nonsocial Odor, and Novel Room conditions (Empty-Session A vs. Empty-Session B: F(1,72) = 2.15, p = 0.88; Nonsocial Odor-Session A vs. Nonsocial Odor-Session B: F(1,72) = 0.03, p = 1; Novel Room-Session A vs. Novel Room-Session B: F(1,72) = 3.20, p = 0.47).These results show that our collection of ethologically relevant stimuli (social interaction, social odor, and fox odor) were sufficiently salient to alter rats’ exploratory behavioral patterns on the track yet were not associated with significant remapping.

### DS1s were differentially modulated by social stimuli

Although nonspatial cues, including social stimuli, did not induce significant remapping in DG place cells, it remained possible that characteristic features of the DG LFP signaled environmental changes. Dentate spikes are population events within the DG LFP that are thought to originate from excitatory entorhinal cortex inputs (Bragin et al., 1995; Dvorak et al., 2021). Dentate spikes are typically split into two types, DS1/L and DS2/M, based on their CSD profiles (but see Tarcsay et al. 2025). DS1 is associated with a current sink (presumed inward current) in the outer molecular layer (OML), whereas DS2 is associated with a current sink in the middle molecular layer (MML) (Bragin et al., 1995; Dvorak et al., 2021; Farrell et al., 2024; McHugh et al., 2024). The DG OML is targeted by excitatory projections from LEC, whereas the DG MML is targeted by excitatory projections from MEC (Amaral, Scharfman, and Lavenex, 2007; Witter, 2007). Thus, DS1s and DS2s are thought to be triggered by LEC and MEC, respectively.

If LEC provides nonspatial, contextual information during DS1s, then a plausible hypothesis is that DS1 rates would increase during presentation of nonspatial stimuli. To test this hypothesis, we assessed the extent to which DS rates changed during exploration of nonspatial stimuli (Fig. 6). We observed a significant 3-way interaction effect between condition, session, and dentate spike type on DS rate (F(5,116) = 2.49, p = 0.035), indicating that dentate spike rates for DS1 and DS2 were differentially affected by condition and session. We next assessed DS1 and DS2 rates separately by reducing the full statistical model into DS1- or DS2-specific models. When the model was fit using only DS1 rates, we still observed a significant interaction between condition and session (F(5,58) = 1.45 x 10^-10^). Fig. 6A shows, for an example rat, the five-second LFP epochs with the maximal rate of DS1s for each condition from session B (i.e., the session in which experimental stimuli were presented). Higher rates of DS1 events were apparent in session B of the Social and Social Odor conditions (Fig. 6B). Indeed, post-hoc comparisons revealed that DS1 rates were significantly higher in session B than in session A of the Social (two-tailed Wald F-test for Social-Session A vs. Social-Session B: F(1,58) = 72.46, p = 5.12 x 10^-11^), Social Odor (Social Odor-Session A vs. Social Odor-Session B: F(1,58) = 57.66, p = 1.77 x 10^-9^), and Fox Odor conditions (Fox Odor-Session A vs. Fox Odor-Session B: F(1,58) = 9.28, p = 0.02). DS1 rates were not different between sessions in the Empty control (Empty-Session A vs. Empty-Session B: F(1,58) = 2.07, p = 0.94) or Nonsocial Odor conditions (Nonsocial Odor-Session A vs. Nonsocial Odor-Session B: F(1,58) = 1.13, p = 1). Interestingly, DS1 rates were significantly lower in the novel room compared to the familiar room (Novel Room-Session A vs. Novel Room-Session B: F(1,58) = 25.86, p = 2.47 x 10^-5^). Finally, to assess whether the Social Odor condition was associated with more DS1s than the Fox Odor condition, a further reduced statistical model was fit using data from Social Odor and Fox Odor conditions only. A significant interaction effect between condition and session was observed (F(1,24) = 5.49, p = 0.028), indicating DS1s were more prevalent in the presence of the social odor than the fox odor. These results suggest that the presentation of ethologically relevant stimuli in an environment may be associated with varying levels of increased input from LEC, reflected by higher rates of DS1 events. In a new spatial location, however, MEC input may take precedence over LEC input, perhaps reflected by a relatively lower rate of DS1 events in a novel room.

If DS2s reflect spatial information transmitted by MEC inputs to the DG, DS2 rates may be expected to increase during exploration of a novel location but not during exploration of nonspatial stimuli. To test this hypothesis, we compared DS2 rates across sessions and conditions (Fig. 7). Example LFP epochs associated with maximal rates of DS2 events for each condition from session B for an example rat are shown in Fig. 7A. Rates of DS2 events appeared similar across conditions and sessions (Fig. 7B). Indeed, the DS2-specific GLMM showed no significant interaction effect between condition and session (F(5,58) = 1.87, p = 0.11). These results suggest that the occurrence of DS2s is not significantly influenced by the presence of ethologically relevant stimuli or novel spatial locations.

We next hypothesized that active exploration of the ethologically relevant stimuli may have elicited the increased DS1 rates. To assess how DS1 rates changed across track positions for each condition and session, we created rate maps for DS1 events (Fig. 8A-F; see *Dentate spike analyses* section of Materials and Methods). Consistent with our hypothesis, we observed a significant 3-way interaction effect between condition, session, and track zone type on DS1 rate (F(10,174) = 3.80, p = 1.22 x 10^-4^), indicating that the observed condition-specific changes in DS1 rates did not occur uniformly across all track positions between sessions. To investigate DS1 rates within each track zone, the full statistical model was reduced into 3 zone-specific models. The Stimulus Zone model was associated with an interaction effect between condition and session (F(5,58) = 4.80, p = 9.71 x 10^-4^) on DS1 rate. Within the Stimulus Zone, DS1 rates were increased in session B compared to session A for the Social (post-hoc two-tailed Wald F-tests: Social-Session A vs. Social-Session B: F(1,58) = 20.08, p = 2.13 x 10^-4^) and Social Odor (Social Odor-Session A vs. Social Odor-Session B: F(1,58) = 19.69, p = 2.48 x 10^-4^) conditions. On the other hand, there were no significant changes in DS1 rates within the Stimulus Zone between sessions A and B for the Empty control (Empty-Session A vs. Empty-Session B: F(1,58) = 0.02, p = 1), Nonsocial Odor (Nonsocial Odor-Session A vs. Nonsocial Odor-Session B: F(1,58) = 0.54, p = 1), Fox Odor (Fox Odor-Session A vs. Fox Odor-Session B: F(1,58) = 0.34, p = 1), and Novel Room (Novel Room-Session A vs. Novel Room-Session B: F(1,58) = 2.02, p = 0.96) conditions. Neither of the other two zone-specific reduced models were associated with a significant interaction effect between condition and session on DS1 rate (Reward Zone: F(5,58) = 2.27, p = 0.059; Neither Zone: F(5,58) = 1.00, p = 0.425), indicating that DS1 rates in locations outside of the Stimulus Zone did not selectively increase between sessions for any particular conditions. Overall, these results support the conclusion that active exploration of social stimuli specifically increased the rate of DS1 events.

### DG place cell activity during dentate spikes

In an attempt to bridge our place cell and dentate spike findings, we investigated the firing of DG place cells during dentate spikes across conditions and sessions. Dentate spikes are associated with granule cell, mossy cell, and interneuron activation (Bragin et al., 1995; Penttonen et al., 1997; Senzai and Buzsáki, 2017; Dvorak et al., 2021; Farrell et al., 2024; McHugh et al., 2024). DS2s have been reported to activate more DG cells than DS1s (Farrell et al., 2024; McHugh et al., 2024), and DS1s have also been associated with inhibition (Dvorak et al., 2021; McHugh et al., 2024). DG cell firing patterns that occurred during waking have been reported to reactivate during dentate spikes in subsequent rest (McHugh et al., 2024). However, how DG place cells fire during dentate spikes occurring during exploration of spatial environments containing different nonspatial stimuli is unknown. The degree to which DS1s and DS2s modulated DG place cell firing is shown in Figs. 9 and S4, respectively. Firing rates during DS1s and DS2s were not significantly altered by presentation of various nonspatial stimuli or exploration of a novel spatial environment (no significant interaction effect between condition, session, and dentate spike type on place cell firing rate modulation (F(5,468) = 0.27, p = 0.93)). Furthermore, DS1- and DS2-associated firing rates did not selectively differ across sessions for any of the experimental conditions (no significant condition by session interaction effects in reduced DS1- and DS2-specific models, DS1: F(5,234) = 0.50, p = 0.78; DS2: F(5,234) = 0.68, p = 0.64). Together, these results suggest that DS1- and DS2-associated DG place cell firing is not significantly affected by the presentation of nonspatial stimuli of varying ethological relevance or exploration of a novel spatial environment.

## Discussion

We investigated DG activity at neuronal and network levels in rats exploring a familiar spatial environment in which nonspatial stimuli varied or a novel spatial environment. We observed no significant changes in the spatial firing patterns of DG place cells, and no significant firing rate changes in place cells with stable place field locations, when nonspatial social and olfactory stimuli were presented. In contrast, significant changes in DG place cells’ spatial firing patterns were observed in a novel environment (Figs. 2-3). At the network level, DS1 rates increased when ethologically relevant stimuli, including social stimuli, were presented but decreased in a novel environment (Fig. 6). Furthermore, DS1 rates were specifically increased around the location where social stimuli were presented (Fig. 8). In contrast, DS2 rates did not differ across experimental conditions (Fig. 7). Finally, environmental changes did not affect DG place cell firing during DS1s (Fig. 9) or DS2s (Fig. S4). Together, these results reveal a novel DG response to social stimuli at the network level and suggest that the DG utilizes different types of processing for spatial and nonspatial information.

In general, statistically significant remapping indicates that changes in place cell firing patterns occurred across a sufficient number of cells to constitute a population-level response to an environmental manipulation (Colgin, Moser, and Moser, 2010). In this study, nonsignificant remapping to nonspatial stimuli suggests that the stimuli used elicited negligible or inconsistent changes in DG place cell firing patterns. However, as can be seen in Figs. 3-4, a minority of cells may have changed their firing patterns when nonspatial stimuli were presented. In the DG, changes in the firing patterns of a single granule cell could initiate downstream changes through the modulation of “detonator” mossy fiber synapses (McNaughton and Morris, 1987; Treves and Rolls, 1992). It is possible that the minority of cells that appeared to change their firing patterns correspond to cue cells that code nonspatial stimuli (Tuncdemir et al., 2022). However, as the spatial location of our nonspatial stimuli was held constant, we cannot rule out the possibility of feature-in-place responses (O’Keefe and Krupic, 2021). It is also possible that the subgroup of cells that changed their firing patterns during presentation of social stimuli correspond to granule cells that project directly to CA2 (Kohara et al., 2014; Llorens-Martin et al., 2015; Laham et al., 2024). Social specificity in the remapping of CA2 place cells (Alexander et al., 2016; Robson et al., 2025) may be conferred by CA2’s enriched expression of receptors for oxytocin, a social neuropeptide (Donahue et al., 2025). Mossy cells and DG interneurons also express oxytocin receptors (Harden and Frazier, 2016; Raam et al., 2017; Hung et al., 2023) and are responsible, in part, for shaping the activity of granule cells. Future studies could causally test whether oxytocin receptor activation induces changes in firing patterns of CA2 or DG place cells when social stimuli are presented.

Due to the sparsity of granule cell firing and the relatively low number of mossy cells, recording a large number of active place cells of each cell type was difficult. To increase total cell yield, we combined potential granule and mossy cells into a single class of DG place cells for remapping analyses. Differences in remapping dynamics between granule and mossy cells have been described previously (Goodsmith et al., 2017; Senzai and Buzsáki, 2017; Kim et al., 2023). As such, remapping in a particular DG cell type in our study may have been obfuscated by differential remapping between granule cells and mossy cells. Nonspatial remapping should be further investigated in each cell type separately in the future.

The LEC transmits nonsocial olfactory information to the hippocampus during olfactory learning (Igarashi et al., 2014; Woods et al., 2020). Also, stimulation of the lateral olfactory tract produces current sinks in the DG OML that are similar to DS1s (Liu and Bilkey, 1997), and dentate spikes are entrained by breathing rate (Karalis and Sirota, 2022). With these findings in mind, we predicted increased DS1 rates during presentation of all olfactory stimuli, under the assumption that DS1s reflect sensory information flowing from LEC to DG. Ethologically relevant stimuli-specific increases in DS1 rate, which were strongest for social stimuli, were therefore unexpected (Figs. 6, 8). LEC inputs to CA2 have also shown social specificity. Exploration of familiar and novel conspecifics, but not novel objects, increases activation of LEC inputs to CA2, and the LEC-to-CA2 projection is necessary for social memory (Lopez-Rojas et al., 2022). However, the contribution of the DG to social memory is disputed (Chiang et al., 2018; Lopez-Rojas et al., 2022; Raam et al., 2017; Leung et al., 2018), and thus it is unclear how DS1s would increase selectively in the presence of social stimuli. It has been suggested that LEC filters out incoming non-salient sensory information and projects only relevant sensory information to the hippocampus (Knierim, Neunuebel, and Deshmukh, 2014). This hypothesis could explain increased DS1 rates in our ethologically relevant stimuli conditions since behavioral exploration data suggested that these stimuli were the most salient (Fig. 5). It should be noted, however, that fox urine presentation surprisingly elicited increased stimulus exploration instead of aversion. It could be that aversive behavior relies on direct contact with the urine sample (Fendt, 2006; Wernecke et al., 2015), which did not occur in this study. The ventral DG is important for anxiety behaviors (Weeden et al., 2015), and anxiety signals may have occurred in the presence of fox odor, regardless of aversion. Thus, it would be interesting to test whether presentation of fox urine modulates the firing of ventral DG place cells during DS1s. Increased DS1 rates during ethologically relevant stimuli presentation, and decreased rates during novel room exploration (Fig. 6), suggest that the influence of LEC inputs on the DG depends on the degree of environmental change. In a familiar environment, a baseline level of DS1s may increase with the addition of a salient nonspatial stimulus. When the spatial environment completely changes to a different spatial location, however, spatially modulated MEC inputs may dominate over nonspatial LEC inputs, resulting in fewer DS1s. If DS2s carry spatial information, and DS1s transmit nonspatial information including olfactory signals, then one may expect DS2 rates to increase in a novel room. However, such an increase in DS2 rates was not observed (Fig. 7). Recent research showed that DS2s can be elicited by loud tones and are associated with CA1 place cell firing that codes current position (Farrell et al., 2024). These findings suggest that DS2s do not simply convey spatial information but rather signal a “miniarousal” state that orients an animal to its current environment (Farrell et al., 2024; Farrell and Soltesz, 2025).

Current literature indicates that DS1s and DS2s occur at different rates. Rats generally display more DS1s than DS2s (compare overall rates in Figs. 6B and 7B; see also Bragin et al., 1995; Paleologos et al., 2025), but studies in mice report the opposite relationship (Dvorak et al., 2021; Farrell et al., 2024; McHugh et al., 2024). A recent mouse study has suggested that differences in the rates of dentate spike types depend on the behavioral state of the animal. DS2s occur more frequently during awake rest and slow-wave sleep, but not running behavior, compared to DS1s (Tarcsay et al., 2025). In the present study, dentate spikes were detected during active behavioral sessions in which rats ran laps on a circular track and paused to investigate experimental stimuli. Thus, differences across studies may also reflect differences in behavior.

Computational modeling suggests that LEC and MEC activity is necessary for DG rate remapping (Rennó-Costa, Lisman, and Verschure, 2010), and rate remapping in CA3 place cells has been shown to rely on LEC input (Lu et al., 2013). Therefore, an exciting hypothesis is that the LEC may contribute to remapping by modulating the firing of DG place cells during DS1s. However, in this study, DG place cells did not show significant firing rate changes during DS1s or DS2s between sessions A and B of any condition (Figs. 9, S4). This finding should be interpreted carefully as we cannot conclude DS1s are not related to rate remapping since we did not observe significant rate remapping in any condition. Significant global remapping was observed, however, in the Novel Room condition without significant firing rate changes during DS1s or DS2s in the novel room. Nevertheless, it remains possible that firing rates during DS1s may be altered in situations in which cells exhibit significant rate remapping. Another possibility is that DS1s modulate the firing of cue cells but not place cells. To clarify the effects of DS1-related inputs on place cells, an important future experiment would involve simultaneous intracellular recordings of DG place cells (Zhang et al., 2020) and extracellular recordings of dentate spikes.

The current study suggests a novel nonspatial coding operation within the DG. Social stimuli and predator odor increased DS1 rates but did not consistently alter DG place cell firing patterns. Further work is necessary to elucidate how dentate spikes and place cells interact during ethologically relevant behaviors.

## Acknowledgments

This research was supported by a National Institutes of Health Award R01MH131317 (to L.L.C.) and a University of Texas at Austin Graduate Continuing Fellowship (to P.G.D.). The authors thank Misty Hill and Jessie Goins for technical assistance and thank Jesus Jimenez and Michael Tsimberg for manually scoring dentate spikes. The authors also thank Jayanth Taranath for valuable assistance in setting up the data acquisition system, providing some of the MATLAB code used for data processing, and valuable discussions. The authors thank Matthew Hersh from the Department of Statistics and Data Sciences at The University of Texas at Austin for statistical consulting. The authors acknowledge the Texas Advanced Computing Center (TACC) at The University of Texas at Austin for providing data storage resources that have contributed to the research described within this article. URL: http://www.tacc.utexas.edu

## Author contributions

PGD designed research, performed research, analyzed data, and wrote the paper; LLC designed research and wrote the paper.

## References

Alexander GM, Farris S, Pirone JR, Zheng C, Colgin LL, Dudek SM (2016) Social and novel contexts modify hippocampal CA2 representations of space. Nat Commun 7:10300.

Allegra M, Posani L, Gómez-Ocádiz R, Schmidt-Hieber C (2020) Differential Relation between Neuronal and Behavioral Discrimination during Hippocampal Memory Encoding. Neuron 108:1103–1112.e6.

Amaral DG, Scharfman HE, Lavenex P (2007) The dentate gyrus: fundamental neuroanatomical organization (dentate gyrus for dummies). Prog Brain Res 163:3–22.

Borzello M, Ramirez S, Treves A, Lee I, Scharfman H, Stark C, Knierim JJ, Rangel LM (2023) Assessments of dentate gyrus function: discoveries and debates. Nat Rev Neurosci 24:502–517.

Buzsáki G, Mizuseki K (2014) The log-dynamic brain: how skewed distributions affect network operations. Nat Rev Neurosci 15:264–278

Bragin A, Jandó G, Nádasdy Z, van Landeghem M, Buzsáki G (1995) Dentate EEG spikes and associated interneuronal population bursts in the hippocampal hilar region of the rat. J Neurophysiol 73:1691–1705.

Chiang MC, Huang AJY, Wintzer ME, Ohshima T, McHugh TJ (2018) A role for CA3 in social recognition memory. Behav Brain Res 354:22–30.

Colgin LL, Moser EI, Moser MB (2008) Understanding memory through hippocampal remapping. Trends Neurosci 31:469–477.

Donahue MM, Robson E, Marron AM, Fernandez EJ, Hill M, Mably AJ, Trimper JB, Brager DH, Colgin LL (2025) CA2 neurons show abnormal responses to social stimuli in a rat model of Fragile X syndrome. bioRxiv 2025.09.10.674743. 10.1101/2025.09.10.674743

Dudek SM, Alexander GM, Farris S (2016) Rediscovering area CA2: unique properties and functions. Nat Rev Neurosci 17:89–102.

Dvorak D, Chung A, Park EH, Fenton AA (2021) Dentate spikes and external control of hippocampal function. Cell Rep 36:109497.

Fabre JMJ, van Beest EH, Peters AJ, Carandini M, Harris KD (2023) Bombcell: automated curation and cell classification of spike-sorted electrophysiology data (1.0.0). Zenodo. 10.5281/zenodo.8172822

Farrell JS, Hwaun E, Dudok B, Soltesz I (2024) Neural and behavioural state switching during hippocampal dentate spikes. Nature 628:590–595.

Farrell JS, Soltesz I (2025) Noncanonical circuits, states, and computations of the hippocampus. Science 389, eadv4420.

Fendt M (2006) Exposure to urine of canids and felids, but not of herbivores, induces defensive behavior in laboratory rats. J Chem Ecol, 32:2617–2627.

Fernández-Ruiz A, Oliva A, Soula M, Rocha-Almeida F, Nagy GA, Martin-Vazquez G, Buzsáki G (2021) Gamma rhythm communication between entorhinal cortex and dentate gyrus neuronal assemblies. Science 372:6537, eabf3119.

GoodSmith D, Chen X, Wang C, Kim SH, Song H, Burgalossi A, Christian KM, Knierim JJ (2017) Spatial Representations of Granule Cells and Mossy Cells of the Dentate Gyrus. Neuron 93:677–690.e5.

GoodSmith D, Lee H, Neunuebel JP, Song H, Knierim JJ (2019) Dentate Gyrus Mossy Cells Share a Role in Pattern Separation with Dentate Granule Cells and Proximal CA3 Pyramidal Cells. J Neurosci 39:9570–9584.

Harden SW, Frazier CJ (2016) Oxytocin depolarizes fast-spiking hilar interneurons and induces GABA release onto mossy cells of the rat dentate gyrus. Hippocampus 26:1124–1139.

Hargreaves EL, Rao G, Lee I, Knierim JJ (2005) Major dissociation between medial and lateral entorhinal input to dorsal hippocampus. Science 308:1792–1994

Hsiao YT, Zheng C, Colgin LL (2016) Slow gamma rhythms in CA3 are entrained by slow gamma activity in the dentate gyrus. J Neurophysiol 116: 2594–2603.

Huang LW, Torelli F, Chen HL, Bartos M (2024) Context and space coding in mossy cell population activity. Cell Rep 43:114386.

Hung YC, Wu YJ, Chien ME, Lin YT, Tsai CF, Hsu KS (2023) Loss of oxytocin receptors in hilar mossy cells impairs social discrimination. Neurobiol Dis 187:106311.

Hwaun E, Colgin LL (2019) CA3 place cells that represent a novel waking experience are preferentially reactivated during sharp wave-ripples in subsequent sleep. Hippocampus 29:921–938.

Igarashi KM, Lu L, Colgin LL, Moser MB, Moser EI (2014) Coordination of entorhinal-hippocampal ensemble activity during associative learning. Nature 510:143–147.

Jun JJ, et al. (2017) Fully integrated silicon probes for high-density recording of neural activity. Nature 551:232–236.

Jung D, Kim S, Sariev A, Sharif F, Kim D, Royer S (2019) Dentate granule and mossy cells exhibit distinct spatiotemporal responses to local change in a one-dimensional landscape of visual-tactile cues. Sci Rep 9:9545.

Karalis N, Sirota A (2022) Breathing coordinates cortico-hippocampal dynamics in mice during offline states. Nat Commun 13:467.

Kerr KM, Agster KL, Furtak SC, Burwell RD (2007) Functional neuroanatomy of the parahippocampal region: the lateral and medial entorhinal areas. Hippocampus 17:697–708.

Kim S, Jung D, Royer S (2020) Place cell maps slowly develop via competitive learning and conjunctive coding in the dentate gyrus. Nat Commun 11:4550.

Kim SH, GoodSmith D, Temme SJ, Moriya F, Ming GL, Christian KM, Song H, Knierim JJ (2023) Global remapping in granule cells and mossy cells of the mouse dentate gyrus. Cell Rep 42:112334.

Knierim JJ, Neunuebel JP, Deshmukh SS (2013) Functional correlates of the lateral and medial entorhinal cortex: objects, path integration and local-global reference frames. Philos Trans R Soc Lond B Biol Sci 369:20130369.

Kohara K, Pignatelli M, Rivest AJ, Jung HY, Kitamura T, Suh J, Frank D, Kajikawa K, Mise N, Obata Y, Wickersham IR, Tonegawa S (2014) Cell type-specific genetic and optogenetic tools reveal hippocampal CA2 circuits. Nat Neurosci 17:269–79.

Laham BJ, Gore IR, Brown CJ, Gould E (2024) Adult-born granule cells modulate CA2 network activity during retrieval of developmental memories of the mother. eLife 12:RP90600.

Leung C, Cao F, Nguyen R, Joshi K, Aqrabawi AJ, Xia S, Cortez MA, Snead OC 3rd, Kim JC, Jia Z (2018) Activation of Entorhinal Cortical Projections to the Dentate Gyrus Underlies Social Memory Retrieval. Cell Rep 23:2379–2391.

Leutgeb JK, Leutgeb S, Moser, MB, Moser EI (2007) Pattern separation in the dentate gyrus and CA3 of the hippocampus. Science 315:961–966.

Liu P, Bilkey DK (1997) Parallel involvement of perirhinal and lateral entorhinal cortex in the polysynaptic activation of hippocampus by olfactory inputs. Hippocampus 7:296–306.

Llorens-Martín M, Jurado-Arjona J, Avila J, Hernández F (2015) Novel connection between newborn granule neurons and the hippocampal CA2 field. Exp Neurol 263:285–292.

Lopez-Rojas J, de Solis CA, Leroy F, Kandel ER, Siegelbaum SA (2022) A direct lateral entorhinal cortex to hippocampal CA2 circuit conveys social information required for social memory. Neuron 110:1559–1572.e4.

Lu L, Leutgeb JK, Tsao A, Henriksen EJ, Leutgeb S, Barnes CA, Witter MP, Moser MB, Moser EI (2013) Impaired hippocampal rate coding after lesions of the lateral entorhinal cortex. Nat Neurosci 16:1085–1093.

Mathis A, Mamidanna P, Cury KM, Abe T, Murthy VN, Mathis MW, Bethge M (2018) DeepLabCut: markerless pose estimation of user-defined body parts with deep learning. Nat Neurosci 21:1281–1289.

McHugh SB, Lopes-Dos-Santos V, Castelli M, Gava GP, Thompson SE, Tam SKE, Hartwich K, Perry B, Toth R, Denison T, Sharott A, Dupret D (2024) Offline hippocampal reactivation during dentate spikes supports flexible memory. Neuron 112:3768–3781.e8.

McNaughton BL, Morris RG (1987) Hippocampal synaptic enhancement and information storage within a distributed memory system. Trends Neurosci 10:408–415.

Nokia MS, Gureviciene I, Waselius T, Tanila H, Penttonen M (2017) Hippocampal electrical stimulation disrupts associative learning when targeted at dentate spikes. J Physiol 595:4961–4971.

O’Keefe J (1976) Place units in the hippocampus of the freely moving rat. Exp Neurol 51:78–109

O’Keefe J, Krupic J (2021) Do hippocampal pyramidal cells respond to nonspatial stimuli? Physiol Rev 101:1427–1456.

Pachitariu M, Sridhar S, Pennington J, Stringer C (2024) Spike sorting with Kilosort4. Nat Methods 21:914–921.

Paleologos N, Vöröslakos M, Gonzalez J, Maslarova A, Aykan D, Liu AA, Buzsáki G (2025) Electroanatomy of hippocampal activity patterns: theta, gamma waves, sharp wave-ripples, and dentate spikes. Front Behav Neurosci 19:1685846.

Penttonen M, Kamondi A, Sik A, Acsády L, Buzsáki G (1997) Feed-forward and feed-back activation of the dentate gyrus in vivo during dentate spikes and sharp wave bursts. Hippocampus 7:437–450.

Pettersen KH, Devor A, Ulbert I, Dale AM, Einevoll GT (2006) Current-source density estimation based on inversion of electrostatic forward solution: effects of finite extent of neuronal activity and conductivity discontinuities. J Neurosci Methods 154:116–133.

Raam T, McAvoy KM, Besnard A, Veenema AH, Sahay A (2017) Hippocampal oxytocin receptors are necessary for discrimination of social stimuli. Nat Commun 8:2001.

Rennó-Costa C, Lisman JE, Verschure PF (2010) The mechanism of rate remapping in the dentate gyrus. Neuron 68:1051–1058.

Robson E, Donahue MM, Mably AJ, Demetrovich PG, Hewitt LT, Colgin LL (2025) Social odors drive hippocampal CA2 place cell responses to social stimuli. Prog Neurobiol 245:102708.

Santiago RMM, Lopes-Dos-Santos V, Aery Jones EA, Huang Y, Dupret D, Tort ABL. (2024) Waveform-based classification of dentate spikes. Sci Rep 14:2989.

Senzai Y, Buzsáki G (2017) Physiological Properties and Behavioral Correlates of Hippocampal Granule Cells and Mossy Cells. Neuron 93:691–704.e5.

Skaggs WE, McNaughton BL, Wilson MA, Barnes CA (1996) Theta phase precession in hippocampal neuronal populations and the compression of temporal sequences. Hippocampus 6:149–172.

Steinmetz NA, et al. (2021) Neuropixels 2.0: A miniaturized high-density probe for stable, long-term brain recordings. Science 372, eabf4588.

Sugar J, Moser MB (2019) Episodic memory: Neuronal codes for what, where, and when. Hippocampus 29:1190–1205.

Tarcsay G, Saxena R, Long R, Shobe JL, McNaughton BL, Ewell LA (2025) Dentate spikes comprise a continuum of relative input strength from the lateral and medial entorhinal cortex. bioRxiv 2025.10.27.684857.

Treves A, Rolls ET (1992) Computational constraints suggest the need for two distinct input systems to the hippocampal CA3 network. Hippocampus 2:189–199.

Tuncdemir SN, Grosmark AD, Turi GF, Shank A, Bowler JC, Ordek G, Losonczy A, Hen R, Lacefield CO (2022) Parallel processing of sensory cue and spatial information in the dentate gyrus. Cell Rep 38:110257.

Tuncdemir SN, Grosmark AD, Chung H, Luna VM, Lacefield CO, Losonczy A, Hen R (2023) Adult-born granule cells facilitate remapping of spatial and non-spatial representations in the dentate gyrus. Neuron 111:4024–4039.e7.

Vöröslakos M, Petersen PC, Vöröslakos B, Buzsáki G (2021) Metal microdrive and head cap system for silicon probe recovery in freely moving rodent. eLife 10:e65859.

Weeden CS, Roberts JM, Kamm AM, Kesner RP (2015) The role of the ventral dentate gyrus in anxiety-based behaviors. Neurobiol Learn Mem 118:143–149.

Wernecke KE, Vincenz D, Storsberg S, D’Hanis W, Goldschmidt J, Fendt M (2015) Fox urine exposure induces avoidance behavior in rats and activates the amygdalar olfactory cortex. Behav Brain Res 279:76–81.

Witter MP (2007) The perforant path: projections from the entorhinal cortex to the dentate gyrus. Prog Brain Res 163:43–61.

Woods NI, Stefanini F, Apodaca-Montano DL, Tan IMC, Biane JS, Kheirbek MA (2020) The Dentate Gyrus Classifies Cortical Representations of Learned Stimuli. Neuron 107:173–184.e6.

Yassa MA, Stark CE (2011) Pattern separation in the hippocampus. Trends Neurosci 34:515–25.

Zhang X, Schlögl A, Jonas P (2020) Selective Routing of Spatial Information Flow from Input to Output in Hippocampal Granule Cells. Neuron 107:1212–1225.e7.

