## Supplemental Figures for "Dynamics of Dentate Gyrus Place Cells and Dentate Spikes During Spatial and Nonspatial Changes in Environments"

**Figure S1**. Histological verification of recording locations. Example cresyl violet-stained histological sections of the hippocampus from all 5 rats are shown. Neuropixels probe tracks can be seen in the DG. Rat 452 was implanted with a single shank 1.0 probe. All other rats were implanted with 4-shank 2.0 probes. Rat 451's probe shanks were implanted parallel with the midline instead of perpendicular, so only one track is visible in a coronal section.

**Figure S2**. Automated detection and classification of dentate spikes. A. Left: Confusion matrix of average F1 scores (harmonic mean of precision and recall) for hand-scored (ground truth) versus automated detection of dentate spikes. Automated detectors differed only in the standard deviation (SD) threshold used to identify peaks in the filtered LFP as putative dentate spikes. Right: The 4.0 SD detector was most consistently the best match with ground truth dentate spikes and was selected for subsequent detection. B. Average CSD depth profiles around dentate spikes detected by different methods. C. Principal components analysis was performed on dentate spike peak time CSD profiles (all profiles from one 10-minute session shown). D. Left: K-means clustering on the $1^{st}$ two principal components from C. Right: Average CSD profiles for the two clusters. Note characteristic DS1 (cluster 1) versus DS2 (cluster 2) CSD profiles. E. Left: CSD profiles from C sorted according to cluster identity. Right: The relative dorsoventral positions of the sink/source reversal points were used to classify clusters as either DS1 (cluster 1) or DS2 (cluster 2).

**Figure S3**. Examples of detected and classified dentate spikes. A. DG LFP depth profile recording arranged dorsoventrally (superior blade only). Note the characteristic increase in amplitudes within the DG from channels ~12-25 and sharp deflections reflecting

DS1s (orange triangles) and a DS2 (purple triangle). B. Top: mean LFP depth profile for DS1s (left) and DS2s (right) from an example 10-minute recording. Bottom: mean ± standard error DS1 (left) and DS2 (right) LFP waveform recorded from the hilar channel with the highest dentate spike amplitude for the same example recording.

**Figure S4**. DS2-associated DG place cell firing across sessions and conditions. Same as Fig. 9 but for DS2s.

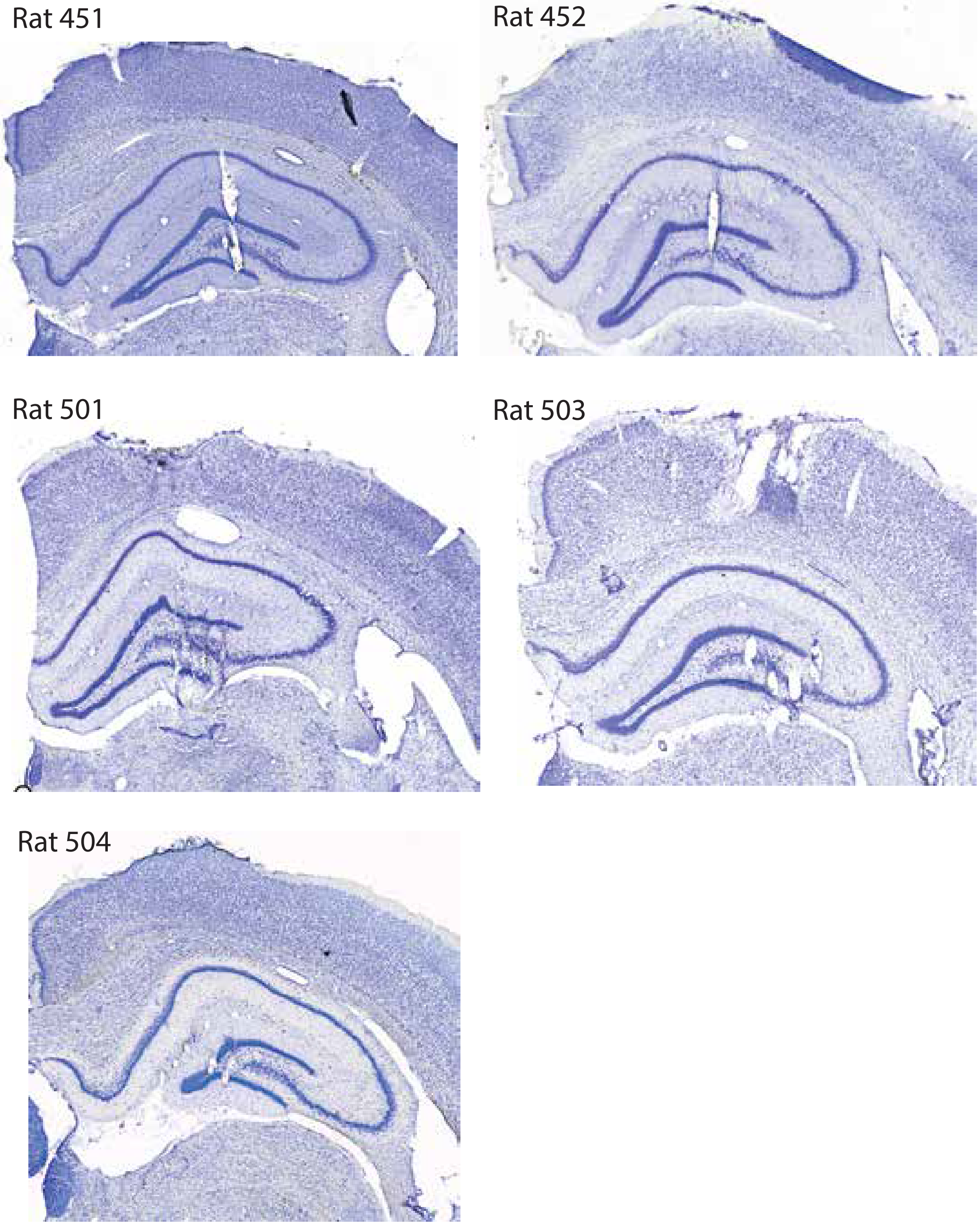

Figure S1

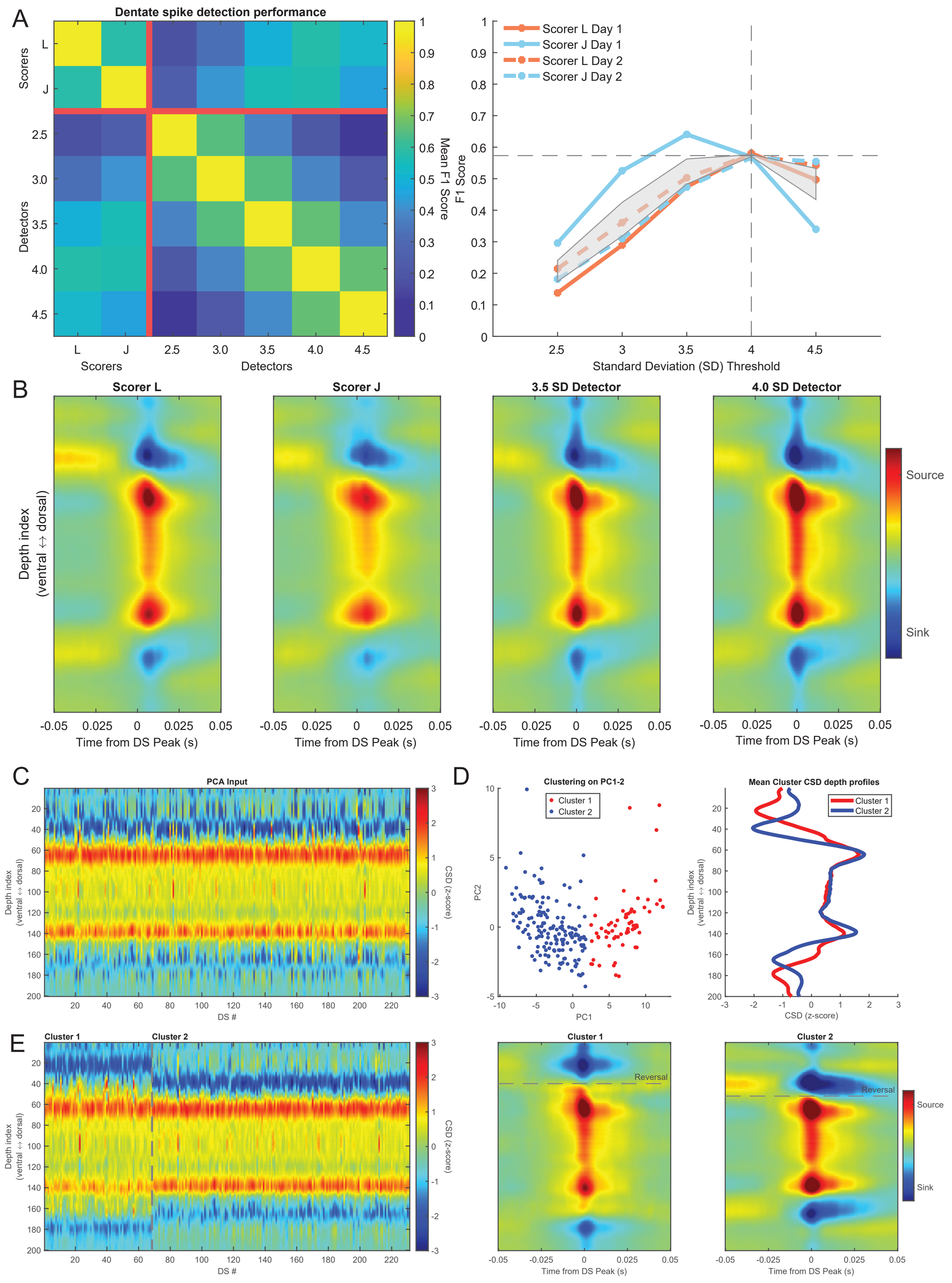
A
Dentate spike detection performance
Scorers
L
J
Detectors
2.5
3.0
3.5
4.0
4.5
Mean F1 Score
F1 Score
Scorer L Day 1
Scorer J Day 1
Scorer L Day 2
Scorer J Day 2
Standard Deviation (SD) Threshold
B
Scorer L
Scorer J
3.5 SD Detector
4.0 SD Detector
Depth index (ventral ↔ dorsal)
Time from DS Peak (s)
Source
Sink
C
PCA Input
Depth index (ventral ↔ dorsal)
DS #
CSD (z-score)
D
Clustering on PC1-2
Cluster 1
Cluster 2
PC1
PC2
Mean Cluster CSD depth profiles
CSD (z-score)
E
Cluster 1
Cluster 2
Reversal
Time from DS Peak (s)

Figure S2

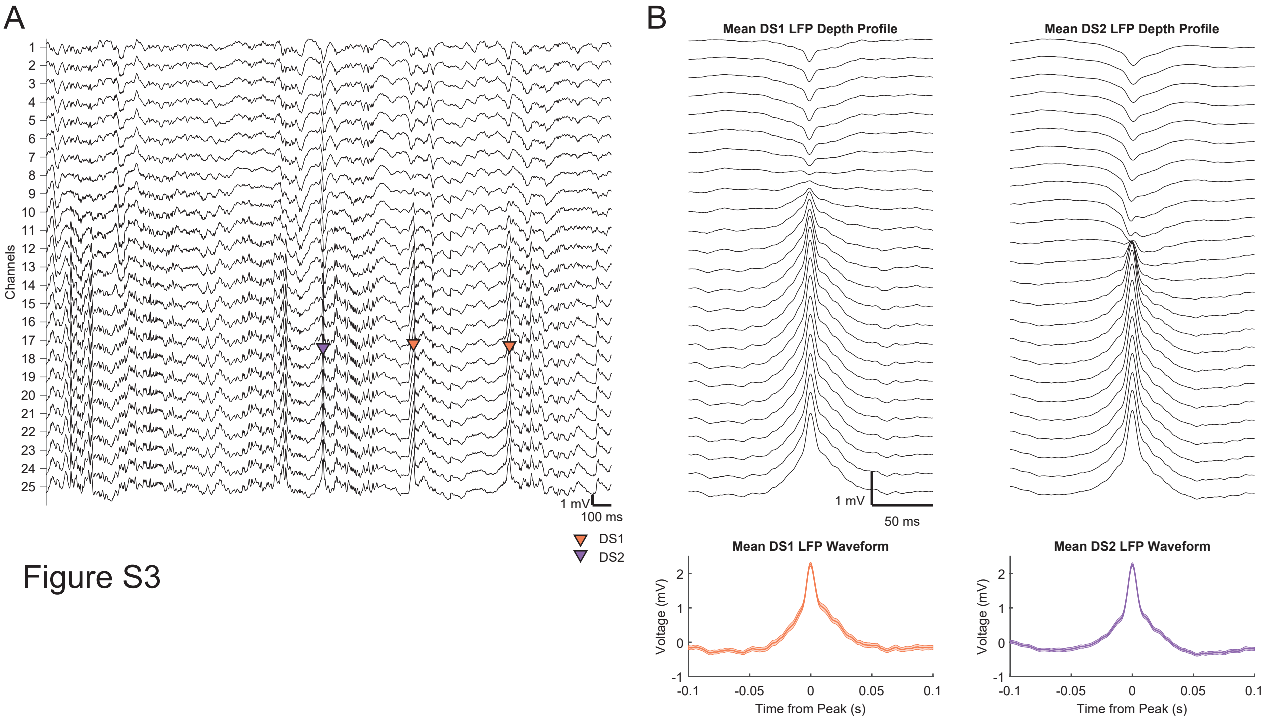
A
B
Channels
1 mV
100 ms
DS1
DS2
Mean DS1 LFP Depth Profile
Mean DS2 LFP Depth Profile
1 mV
50 ms
Mean DS1 LFP Waveform
Mean DS2 LFP Waveform
Voltage (mV)
Time from Peak (s)
-0.1
-0.05
0
0.05
0.1
2
1
0
-1

Figure S3

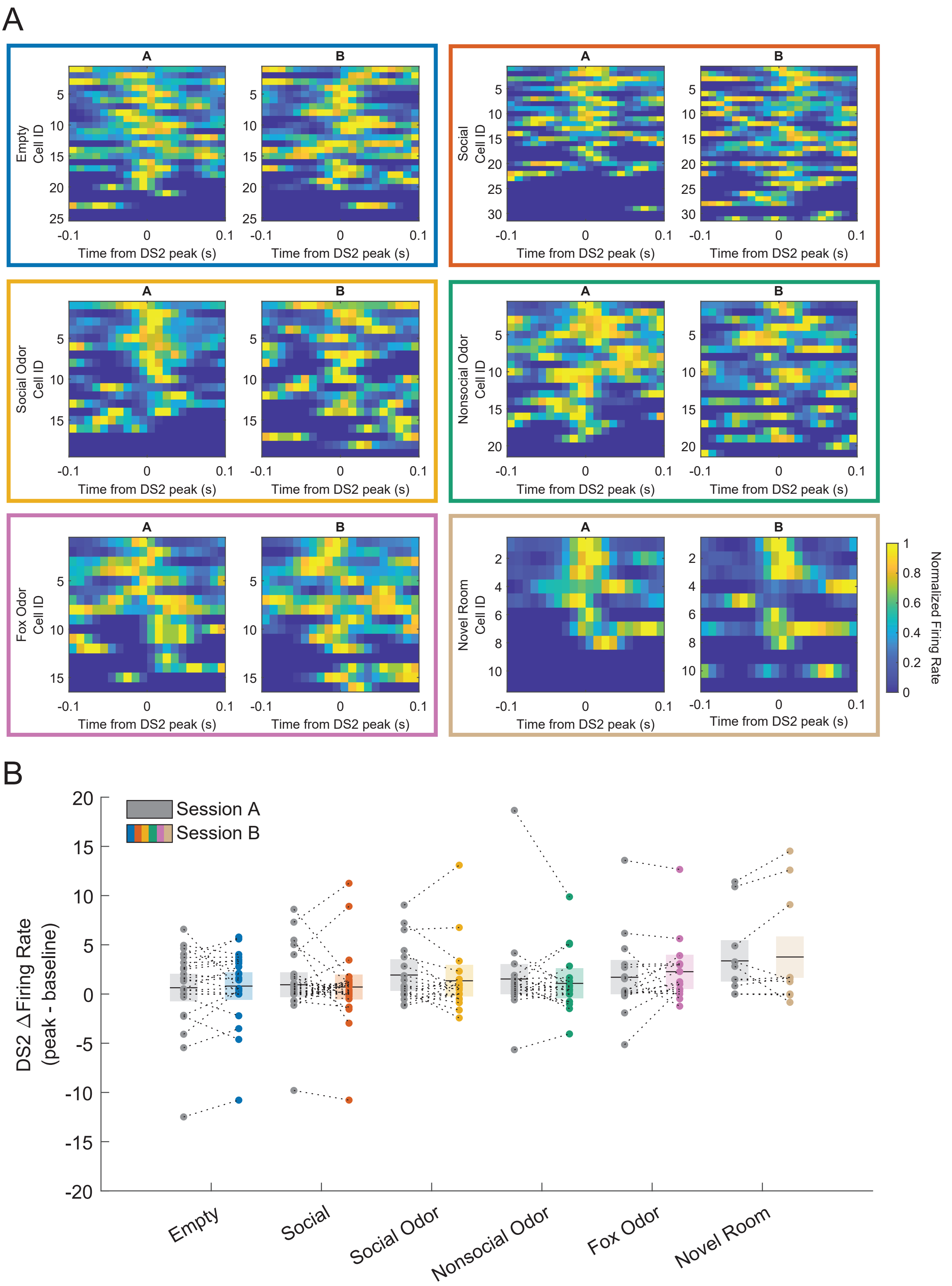
A
A
B
Empty
Cell ID
Time from DS2 peak (s)
Social
Cell ID
Social Odor
Cell ID
Nonsocial Odor
Cell ID
Fox Odor
Cell ID
Novel Room
Cell ID
Normalized Firing Rate
B
Session A
Session B
DS2 ΔFiring Rate
(peak - baseline)
Empty
Social
Social Odor
Nonsocial Odor
Fox Odor
Novel Room

Figure S4
